# Flexible control of representational dynamics in a disinhibition-based model of decision making

**DOI:** 10.1101/2022.04.18.488670

**Authors:** Bo Shen, Kenway Louie, Paul Glimcher

**Affiliations:** Neuroscience Institute, New York University Grossman School of Medicine, New York, NY, United States, 10016; Center for Neural Science, New York University, New York, NY, United States, 10003

**Keywords:** Decision-making circuit, disinhibition, divisive normalization, winner-take-all choice, persistent activity

## Abstract

Current models utilize two primary circuit motifs to replicate neurobiological decision making. Recurrent gain control implements normalization-driven relative value coding, while recurrent excitation and non-selective pooled inhibition together implement winner-take-all (WTA) dynamics. Despite evidence for concurrent valuation and selection computations in individual brain regions, existing models focus selectively on either normalization or WTA dynamics and how both arise in a single circuit architecture is unknown. Here we show that a novel hybrid motif unifies both normalized representation and WTA competition, with dynamic control of circuit state governed by local disinhibition. In addition to capturing empirical psychometric and chronometric data, the model produces persistent activity consistent with working memory. Furthermore, the biological basis of disinhibition provides a simple mechanism for flexible top-down control of network states, enabling the circuit to capture diverse task-dependent neural dynamics. These results suggest a new biologically plausible mechanism for decision making and emphasize the importance of local disinhibition in neural processing.

## Introduction

Two fundamental processes in decision making are value coding and option selection. In the early stage of a decision, value representations serve as integrated decision variables that combine outcome information such as expected gain and probability of realization. Such representations are central to formal decision theories in ecology, economics, and psychology^1^. Consistent with behavioral theories, neural firing rates vary with option values in numerous decision-related brain areas, including the frontal^2–10^ and parietal^10–21^ cortices and basal ganglia^22–25^. Recent research shows that neural value coding is contextual in nature, with the value of a given option represented relative to the value of available alternatives^2,9,15,19,26–30^. Furthermore, this relative value coding employs divisive normalization^9,27,31,32^, a canonical computation prevalent in sensory processing and thought to implement efficient coding principles^33–39^ and temporal adaptation^28,32,36,40–42^.

As the decision process progresses, a common and powerful neural mechanism for categorical choice is winner-take-all (WTA) competition^43,44^. WTA dynamics are widely observed in multiple brain regions: the neural firing rate representing the chosen option or action target increases in concert with selection (often reaching a common activity threshold at choice), while firing rates representing the other unchosen alternatives are suppressed, resulting in a single categorical choice^14–16,19,20,45–50^. The wide prevalence of WTA dynamics in decision-related neural activities suggests that it is a general feature of biological choice.

While valuation and selection processes may occur independently in different brain areas, electrophysiological evidence shows coexisting value coding and WTA signals in prominent decision-related circuits. When decisions are framed as action selection, such integrated representation exists primarily in frontoparietal areas tightly linked to motor action commitment. For example, in the control of eye movements, value and selection dynamics coexist in multiple brain regions including LIP^12,14–17^, the frontal eye fields^3,6,23^, and the superior colliculus^51–55^. In these areas, neural activity initially represents the relevant decision variables, but shifts to encode the selected saccade after a WTA-like interval. Similar activity emerges in parallel circuits controlling arm movements, including the parietal reach region^56–58^, dorsal premotor cortex^2,8,59^, and primary motor cortex^7,8^. Notably, when examined, contextual value coding during a decision typically arises after initial absolute value coding^2,15,29^, consistent with a local normalization process; these dynamics suggest that normalized value coding is not simply inherited from upstream regions and support coexisting within-region normalization and selection computations.

Computational decision models have identified core circuit motifs that produce either normalized value representation or WTA selection (**Fig. 1**). For normalized value representation, dynamic circuit-based models emphasize a crucial role for both lateral and feedback inhibition^29,60^. In the dynamic normalization model (DNM), paired excitatory and inhibitory neurons represent each choice option (**Fig. 1A**); feedforward excitation inputs value information, lateral connectivity mediates contextual interactions, and feedback inhibition drives divisive scaling. This simple differential equation model emphasizes a crucial role of lateral connectivity and feedback inhibition in driving empirically-observed divisive scaling and contextual interactions (**Fig. 1B**).

**Figure 1.**
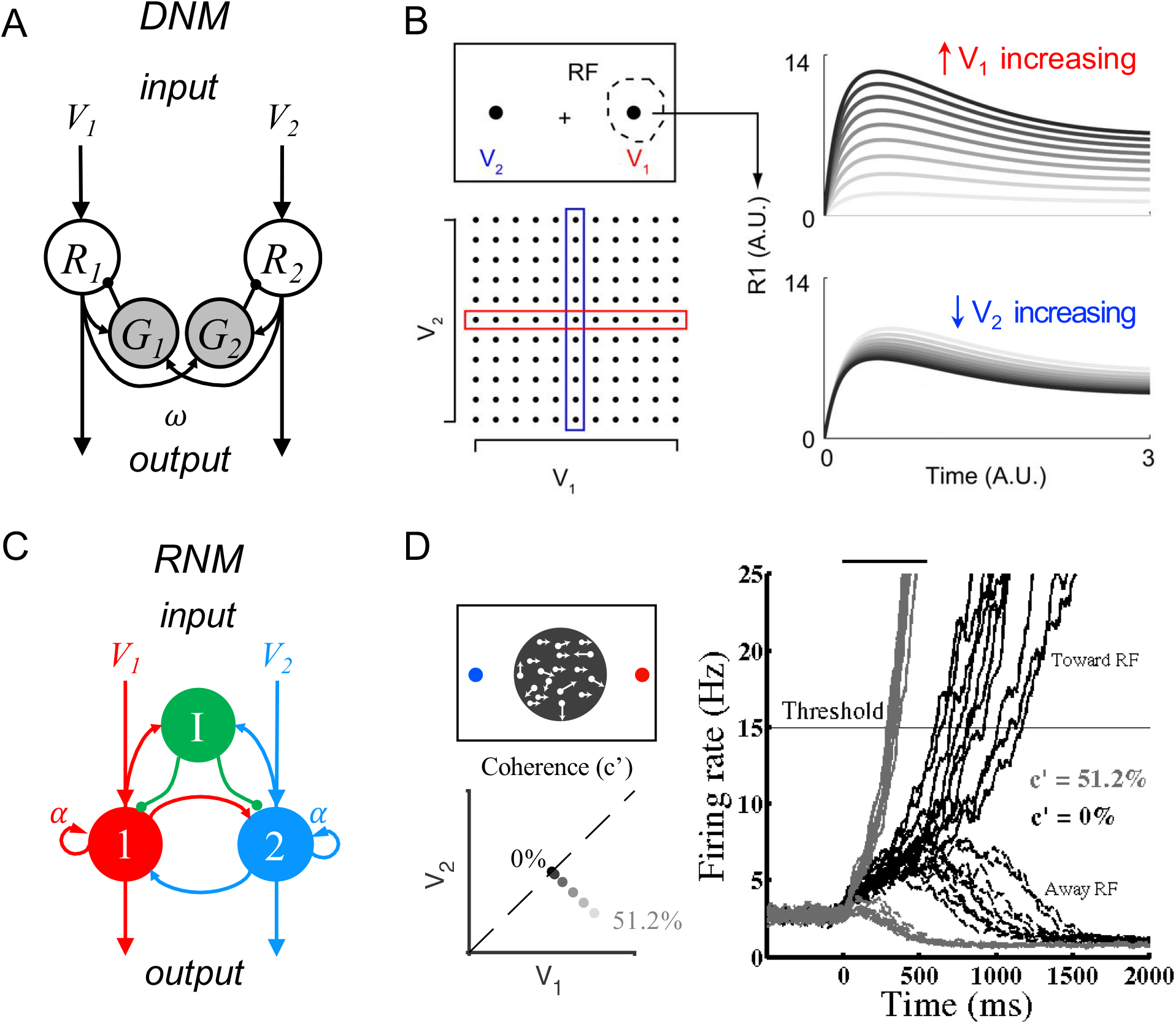
Standard circuit motifs and neural dynamics in existing decision-making models. **A**. Dynamic normalization model (DNM). Each pair of excitatory (*R*) and inhibitory (*G*) units corresponds to an option in the choice set, with *R* receiving value-dependent input *V* and providing output. Lateral interactions implement a cross-option gain control that produces normalized value coding. **B**. DNM predicted dynamics replicate empirical contextual value coding. Example task involves orthogonal manipulation of both option values. *R*_*1*_ activity increases with the direct input value *V*_*1*_ (array framed in red) but is suppressed by the contextual input *V*_*2*_ (array framed in blue), consistent with value normalization. **C**. Recurrent network model (RNM). The network consists of excitatory pools with self-excitation (1 and 2) and a common pool of inhibitory neurons (I). **D**. RNM predicted dynamics generate winner-take-all (WTA) selection. Example task involves motion discrimination of the main direction of a random dot motion stimulus with varying coherence (c’) levels (left). Model activity (right) under two different levels of input coherence (0% and 51.2%) predicts different ramping speeds to the decision threshold, and generates a selection even with equal inputs. Graphics A and B adapted from Louie et al. (2014); graphics C and D adapted from Wong & Wang (2006), Copyright 2006 Society for Neuroscience.

For WTA selection, the predominant class of decision models (recurrent network models, hereafter RNM) propose a central role for recurrent connectivity^61–67^ and non-selective feedback inhibition^43,44^ (**Fig. 1C**). RNMs replicate psychophysical and neurophysiological results in perceptual^49,50,68,69^ and economic^31,70–72^ choices, capturing the complex nonlinear dynamics of empirical neurons (**Fig. 1D**). Furthermore, the competitive nature of the RNM generates attractor states which maintain continued activity even in the absence of stimuli, consistent with persistent spiking activity associated with working memory during delay intervals^48–50,68,73–81^.

Despite the existence of prominent models for normalized valuation and WTA selection, no current model integrates both properties within a single unified circuit. The current DNM cannot capture late-stage choice dynamics because it lacks a mechanism for WTA competition and categorical choice. Similarly, due to the lack of structured lateral inhibition, RNMs typically neither exhibit contextual value coding nor predict contextual choice patterns^82^. Thus, existing DNM and RNM models highlight a need for an integrated decision model that explains both the normalized value coding and option selection seen in the same single brain regions.

To unify these key features of decision-making, we develop and characterize a novel biological circuit consisting of three neuronal types including local disinhibition. At its core, this model hybridizes the architectural features of divisive gain control and recurrent self-excitation used in existing models, but utilizes disinhibition rather than the commonly-assumed pooled inhibition to implement competition for two reasons. First, in contrast to the non-selective or broadly tuned inhibition seen in visual cortex during stimulus representation^83–89^, increasing evidence shows that inhibitory neurons are selective in coding decision variables in frontal cortex^90^, parietal cortex^90,91^, and striatum^92^, suggesting structured inhibition instead of pooled inhibition may play a role in decision circuits. Second, GABAergic inhibitory interneurons exhibit a diversity of subtypes with distinct information processing roles in local circuits^93–97^, including a prominent role for local disinhibition^98–103^. In disinhibitory architectures, vasoactive intestinal peptide (VIP)-expressing interneurons inhibit the neighboring interneurons expressing somatostatin (SST) or parvalbumin (PV) that inhibit dendritic or perisomatic areas in pyramidal neurons, respectively^100–109^ (**Fig. 2C**). Local circuit inputs to VIP neurons suggest that disinhibition may be a key mechanism for generating the mutual competition necessary in option selection and driving selective inhibition. In addition, given the existence of long-range inputs^94,100,102,103,110^ and neuromodulatory inputs^96,97,102,105,111,112^ to VIP neurons, local disinhibition has been proposed to play a particular role in dynamic gating of circuit activity; such gating may be essential in decision circuits underlying flexible behavior, mediating top-down control of network function^100,101,103,105,110,113,114^. We find that a novel hybrid model incorporating recurrent excitation, lateral inhibition, and local disinhibition unifies multiple characteristics of decision activity including normalized value coding, WTA choice, and working memory. These findings suggest that local disinhibition provides a robust, biologically plausible integration of normalization and WTA selection in a single circuit architecture. Moreover, a top-down gating signal operating via this disinhibition enables the model to reproduce decision activity in a range of experimental paradigms with diverse task timing and activity dynamics.

**Figure 2.**
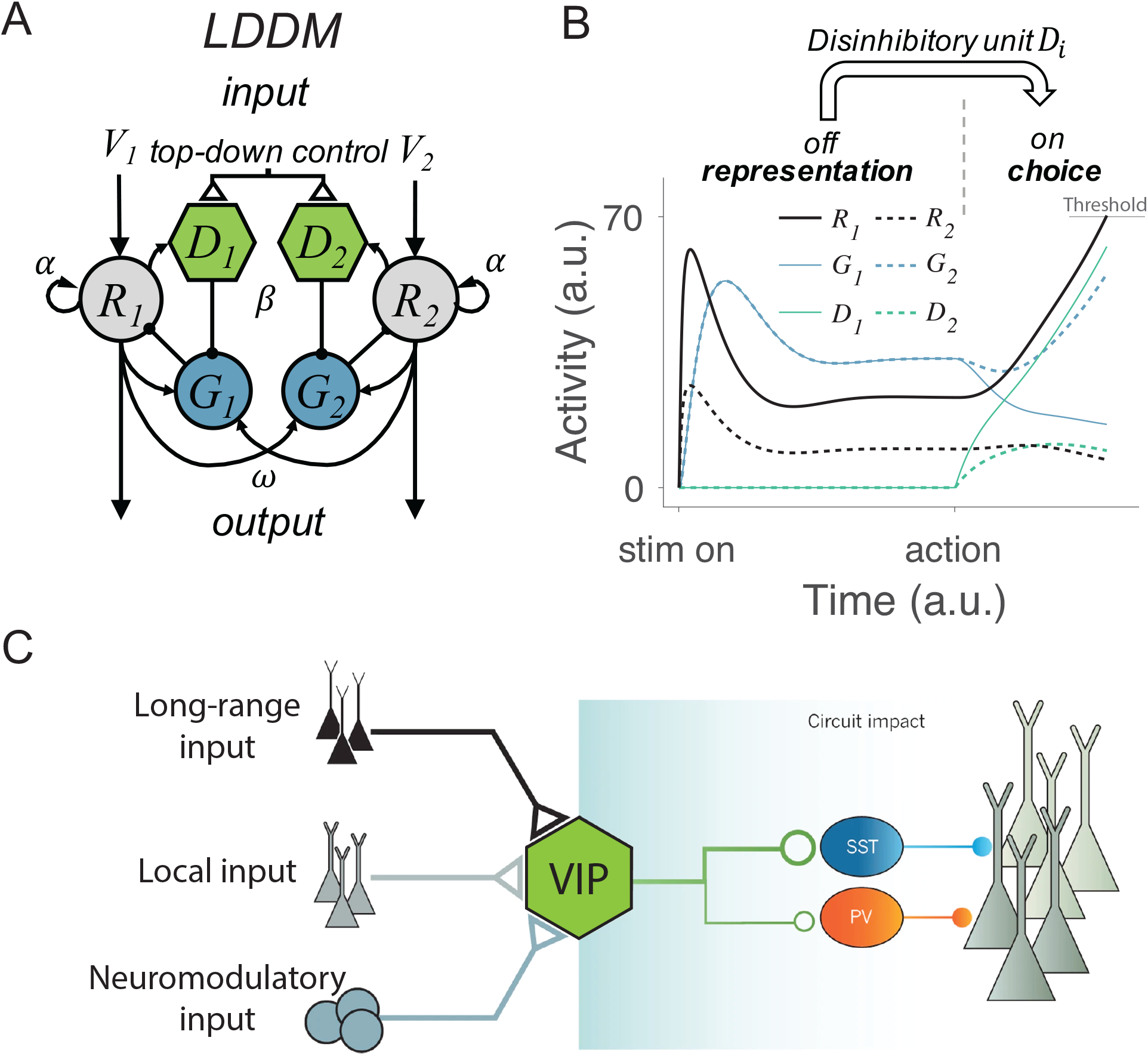
Local disinhibition decision model (LDDM) and its biological plausibility. **A**. LDDM hybridizes DNM and RNM by incorporating a disinhibitory *D* unit to mediate the local disinhibition of the associated excitatory *R* unit; strength of disinhibition is controlled by the parameter *β* presumed via an external top-down control. *V*_*i*_, *α*, and *ω* indicate the corresponding input value to each option, self-excitation of *R* unit, and the coupling weights from *R* to *G* unit, respectively. **B**. The network phase transition between representation and choice under gated disinhibition. With the disinhibitory module silent, the network performs dynamic divisive normalization on *R* units and predicts non-selective inhibition via *G* units; after the disinhibitory module is triggered via an external top-down control signal, the network switches to a winner-take-all competition dynamic and predicts selective inhibition via *G* and *D* units. **C**. Biological basis of disinhibition. Disinhibition provides a mechanism for dynamic gating of circuit states. Vasoactive intestinal peptide (VIP)-expressing interneurons typically inhibit somatostatin (SST) and parvalbumin (PV)-positive interneurons, resulting in a disinhibition of pyramidal neurons. VIP neurons receive local, long-range, and neuromodulatory input, providing different potential mechanisms to modulate local circuit dynamics. Adapted with permission from Kepecs & Fishell (2014).

**Figure 2.**
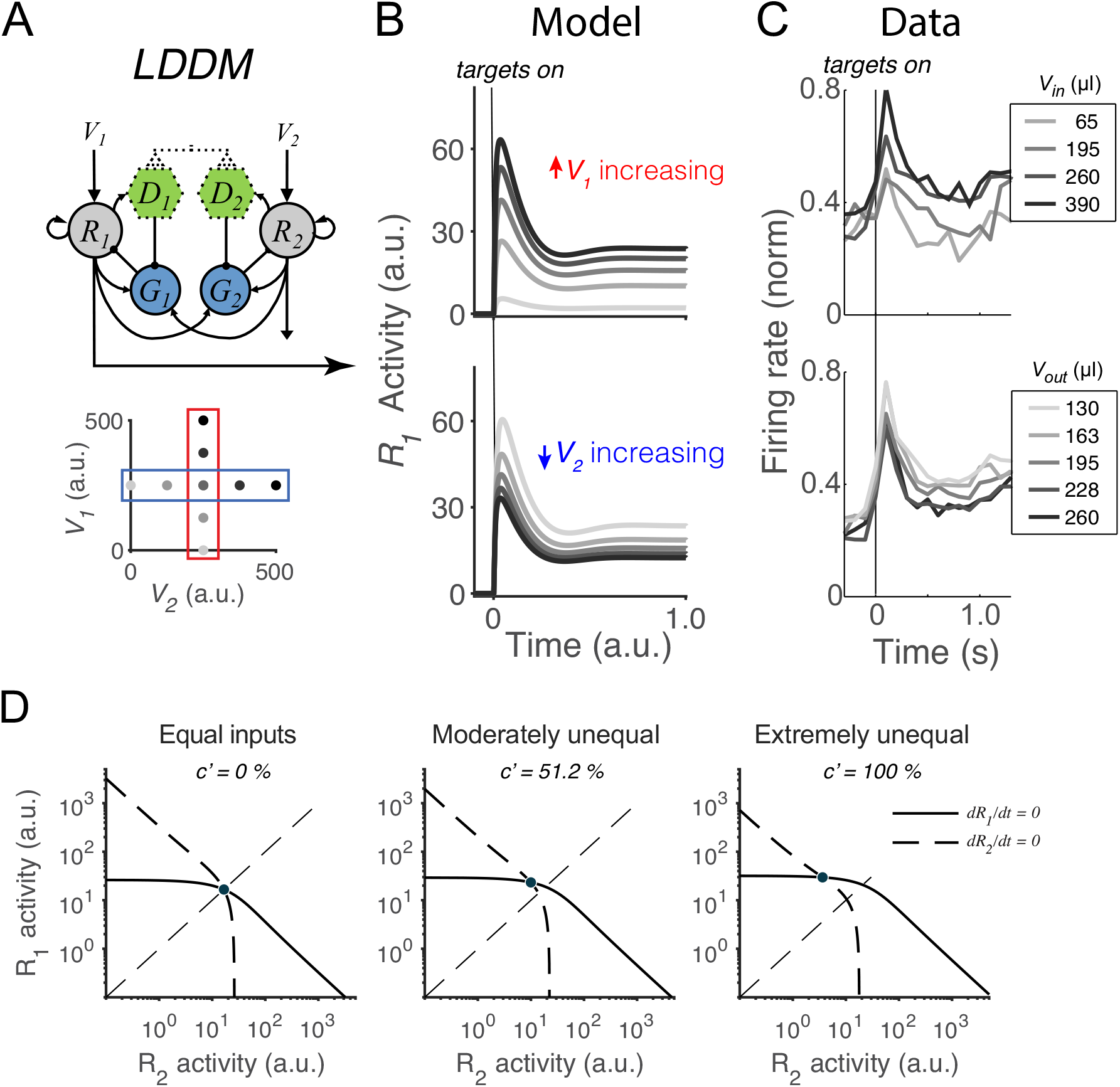
Normalized value coding in the LDDM. **A**. In this example, the LDDM receives a set of two input values with varying *V*_*1*_ (framed in red) and *V*_*2*_ (framed in blue). **B**. Example LDDM dynamics show relative value coding. *R*_*1*_ activity shows a transient peak before a sustained period of coding. Increasing *V*_*1*_ increases *R*_*1*_ activity but increasing *V*_*2*_ decreases *R*_*1*_ activity. **C**. Value coding dynamics recorded in monkey parietal cortex. The model prediction we showed is consistent with the empirical observation. Reprinted from Louie et al. (2011). **D**. Phase-plane analysis of the system under equal (left), weakly unequal (middle), and extremely unequal (right) inputs. The nullclines of *R*_*1*_ (solid) and *R*_*2*_ (dashed) indicating the equilibrium state of the individual units intersect at a unique and stable equilibrium point with divisively normalized coding.

In addition to its ability to capture a broader range of phenomena than existing models, the unique structure of the model offers several other advantages. The model incorporates recent findings from optogenetic studies to provide new information about the cortical microcircuit, employing the known the types and connectivities of inhibitory interneurons to yield observed physiological behavior ^94,96,103,105,109^. It accomplishes this by, unlike prior models, employing the motif of anatomically structured, rather than pooled, inhibition. While structured inhibition is similar to changes of excitatory/inhibitory balance in some ways, the use of this alternative approach allows the model to predict classes of input-selective inhibition that have recently been observed in empirical studies^90–92^, but which cannot be predicted by more traditional motifs. Further, recent empirical studies of fast modulation suggest the importance of structured disinhibition to this process^101,103,110–112,115–118^, a finding well aligned with the model. In summary, the local disinhibition-based decision model (LDDM) offers a new circuit motif that takes account of the most anatomical and physiological data on the nature of structured inhibition which providing a more accurate capture of observed physiological findings during all phases of the decision-making process.

## Results

### Local disinhibition decision model (LDDM)

To develop an integrated circuit model of decision making, we systematically tested a series of models incorporating the core elements of existing models, namely divisive gain control, recurrent excitation, and mutual competition (supplementary **Fig. S1**). This analysis identified *local disinhibition* as the crucial component that can integrate mutual competition and value normalization within the existing circuit architecture of DNM. In the rest of this paper, we focus on the novel disinhibitory hybrid model (local disinhibition decision model, hereafter LDDM).

In the LDDM (**Fig. 2A**), an option-specific disinhibitory *D* unit receives input from its associated excitatory *R* unit and inhibits the inhibitory *G* unit in the local sub-circuit. Biased disinhibition – via different value inputs to option-selective *R* units – can thus introduce an effective asymmetry to global inhibition, generating an unbalanced gain control between local and opponent circuits and leading to WTA competition. In this model, the network shifts from value coding to WTA competition regimes in response to an onset of disinhibition (controlled by the coupling strength between *R* and *D*). With zero or weak *R-D* coupling, the circuit preserves normalized value coding consistent with the DNM; with strong *R-D* coupling, the circuit switches to a state of WTA selection (**Fig. 2B**). Inhibitory units, as a result, dynamically switch from a non-selective response pattern to a selective response pattern (*G* and *D* units in **Fig. 2B**). This flexible onset of disinhibition is modeled after biological findings, which show that activation of disinhibition in cortical circuits arises from exogenous, long-distance projections^100,103,105,113,114^ (**Fig. 2C**). This form of top-down control allows for flexibility in the relative timing of the valuation and selection processes, consistent with neural and behavioral data in different task paradigms (see *Gated disinhibition provides top-down control of choice dynamics*).

Activity dynamics of the LDDM are described by a set of differential equations:

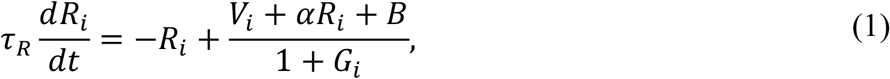

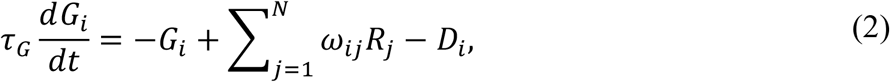

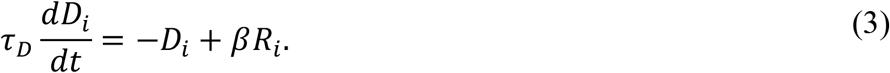

where *i* = 1, …, *N* designates choice alternatives, each of which receive selective input *V*_i_ and non-selective baseline input *B. τ* _R_, *τ* _G_, and *τ* _D_ are the time constants for the *R, G*, and *D* units. The weights *ω*_ij_ represent the coupling strength between excitatory units *R*_j_ and gain control units *G*_*i*_, with each inhibitory unit driven by a weighted sum of its excitatory inputs; the parameter *α* controls the strength of recurrent self-excitation on *R* units. Finally, *β* weights the coupling strength between the excitatory *R*_*i*_ and the disinhibitory *D*_*i*_ units and is presumed to be under external (task-triggered) control.

### Dynamic divisive normalization preserved in the LDDM

We first examine whether the LDDM retains the dynamics of divisively normalized value coding seen in the DNM^29,60^. As discussed above, during initial option evaluation the disinhibitory units are silent (*β* = 0) and the sole difference between the LDDM and the DNM is recurrent excitation (controlled by *α*). Example activity traces in **Fig. 3B** show that the LDDM preserves characteristic early-stage dynamics and contextual modulation seen in both empirical data (**Fig. 3C**) and the original DNM^27,29,60^. Immediately after stimulus onset, *R*_*1*_activities replicate the transient peak observed in a wealth of studies^11,13,15,17,19,27,29,119^. Further, the network settles to equilibrium displaying relative value coding: *R*_*1*_activity increases with *V*_*1*_and decreases with *V*_*2*_ (**Fig. 3B**, *R*_*1*_activity across *V*_*1*_inputs (upper panels) and *V*_*2*_ inputs (bottom panels)), reflecting a contextual representation of value (see Methods for details of parameters used in visualization).

Taking advantage of its simplified mathematical form, we analytically evaluated the LDDM and found that it represents each set of input values (*V*_1_, …, *V*_*N*_) as one unique and stable equilibrium point in its output space (*R*_1_, …, *R*_*N*_) when *β* = 0. **Fig. 3D** shows that the nullclines of *R*_*1*_(solid) and *R*_*2*_ (dashed), which represent the equilibrium state of each individual unit, intersect at a unique and stable equilibrium point regardless of equal or unequal inputs (see *Equilibria and stability analysis of the LDDM* in **Supplementary Results** for the mathematical proof). The steady state of neural activity 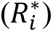 at the equilibrium reflects divisive normalization (Eq. 4), as in the original DNM^29,60^. The only difference at the equilibrium is the negative constant (−*α*) in the denominator introduced by recurrent excitation; this change rescales the activity magnitudes but preserves normalized value coding.

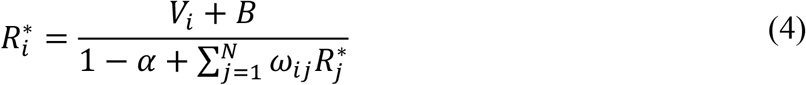

We next verified that the normalized value coding produced by the LDDM cannot be implemented by standard recurrent RNM models. **Fig. 4A** compares the activity of 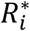 as a function of both value inputs (*V*_*1*_ and *V*_*2*_) in the LDDM (left panel), the original DNM (middle panel), and the RNM (right panel). Both the LDDM and the DNM exhibit 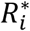 activities (indicated by color) that monotonically increase with input *V*_1_ but decrease with *V*_2_, with a slightly steeper *V*_2_ dependence in the LDDM versus the DNM model depending on the rescaling of *α*. In contrast, strong WTA dynamics in the RNM implement categorical (choice) coding rather than relative value representation, with high or low coding of input values (right panel).

**Figure 4.**
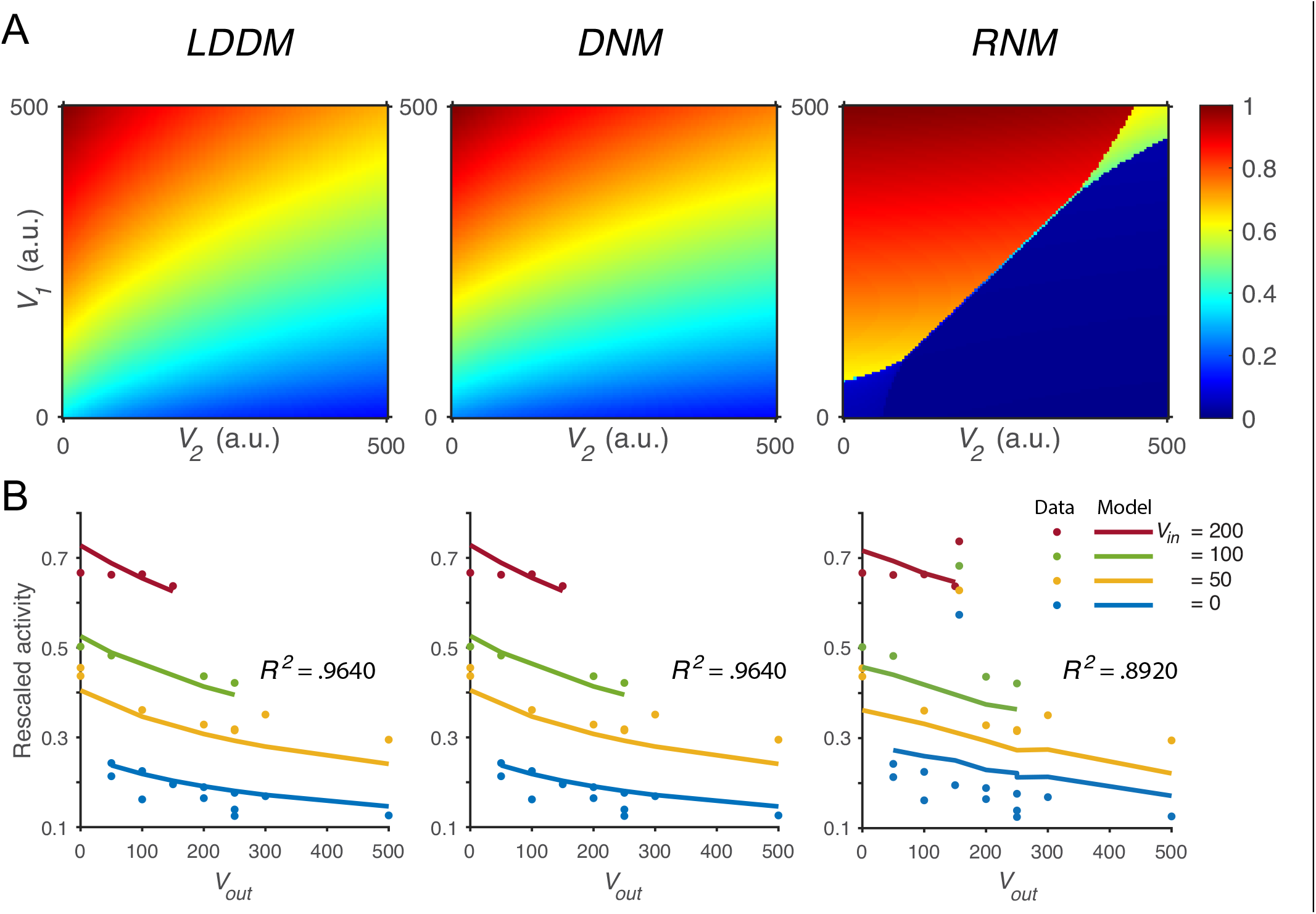
Quantitative comparison of contextual value coding across the LDDM, DNM, and RNM models. **A**. Comparison between the LDDM (left), the DNM (middle), and the RNM (right) in value coding. The LDDM and the DNM show normalized value coding. The neural activity of *R*_*1*_ (indicated by color) increase with the direct input *V*_*1*_ but decrease with the contextual input *V*_*2*_. The LDDM shows slightly stronger contextual modulation than the DNM but qualitatively replicated normalized value coding. The RNM shows a qualitatively different pattern consistent with WTA competition. Within the regime of WTA competition (*V*_*1*_ and *V*_*2*_ within a reasonable scale), *R*_*1*_ activity is high when *V*_*1*_ > *V*_*2*_ and low when *V*_*1*_ < *V*_*2*_. **B**. Fitting the models to a trinary choice dataset shows that the LDDM (left panel) and the DNM (middle panel) precisely capture the neural activities responding to values inside (*V*_*in*_) and outside (*V*_*out*_) of the receptive field. Fitting the RNM to the dataset does not capture the neural activities as well as the LDDM (and DNM) (right panel); furthermore, in this parameter regime the RNM no longer implements WTA selection.

To quantitatively test value normalization, we fit the models to observed firing rates of monkey lateral intraparietal (LIP) neurons under varying reward conditions^27^. In the empirical data (**Fig. 4B**, dots), LIP activity increases with the reward (juice quantity) associated with the target inside the neuronal response field (*V*_*in*_) and decreases with the summed rewards of targets outside the response field (*V*_*out*_). The fitting results show that the DNM precisely captures the rescaled firing rates with only one free parameter (*B =* 70.6 in Eq. 1) (middle panel in **Fig. 4B**, *R*^*2*^ = .9640). Importantly, value normalization is equivalently replicated by the LDDM (with one additional parameter *α*; left panel in **Fig. 4B**, *R*^*2*^ = .9640, *B* = 71.2, *α* = 0). Note that the LDDM reduces to the DNM in this case with *α* = 0; however, a non-zero self-excitation parameter may be important in other scenarios (see *A novel form of persistent activity*).

In contrast to the LDDM (and DNM), fitting the standard RNM with four parameters (see supplementary **Fig. S4** for details) does not capture very well the magnitudes of neural activity as a function of *V*_*in*_ (right panel in **Fig. 4B**) (*R*^2^ = .8920). The curvature of neural activity as a function of *V*_*out*_shows a linear type of lateral inhibition, contrast to the concave curvature predicted by divisive normalization in LDDM (and DNM). Furthermore, fitting the RNM to the data results in a parameter regime that can no longer generate WTA competition; instead, the model predicts mean firing rates in a low-activity regime with maximum value ∼ 3.5 Hz (supplementary **Fig. S4**).

### Local disinhibition drives winner-take-all competition

A key question is whether the LDDM also produces WTA competition. Given the architecture of the LDDM, local disinhibition is hypothesized to break the symmetry between option-specific *R*-*G* sub-circuits, enabling a competitive interaction between sub-circuits. To examine whether this competition produces WTA selection, we simulated model activity in a reaction-time version of a motion discrimination task, a standard perceptual decision-making paradigm in non-human primates^14,19^. The task contains two stages of processing: the pre-motion stage with only the choice targets presented and the motion stage presenting a random-dot motion stimulus. Animals are allowed to select an option, indicating their percept of the main direction of motion, at any time following motion stimulus onset (see timeline, **Fig. 5A**). During the pre-motion stage, we simulated equal value inputs, given the equal prior probability of either target being correct. The simulated pre-motion dynamics replicate the characteristic transient peak observed in both perceptual and economic decision-making tasks^11,15,19,27^. At motion stimulus onset, inputs to the two *R* units are changed according to the task design; because the animals could make their decision at any time in this reaction-time task, disinhibition is assumed to increase when motion inputs begin.

**Figure 5.**
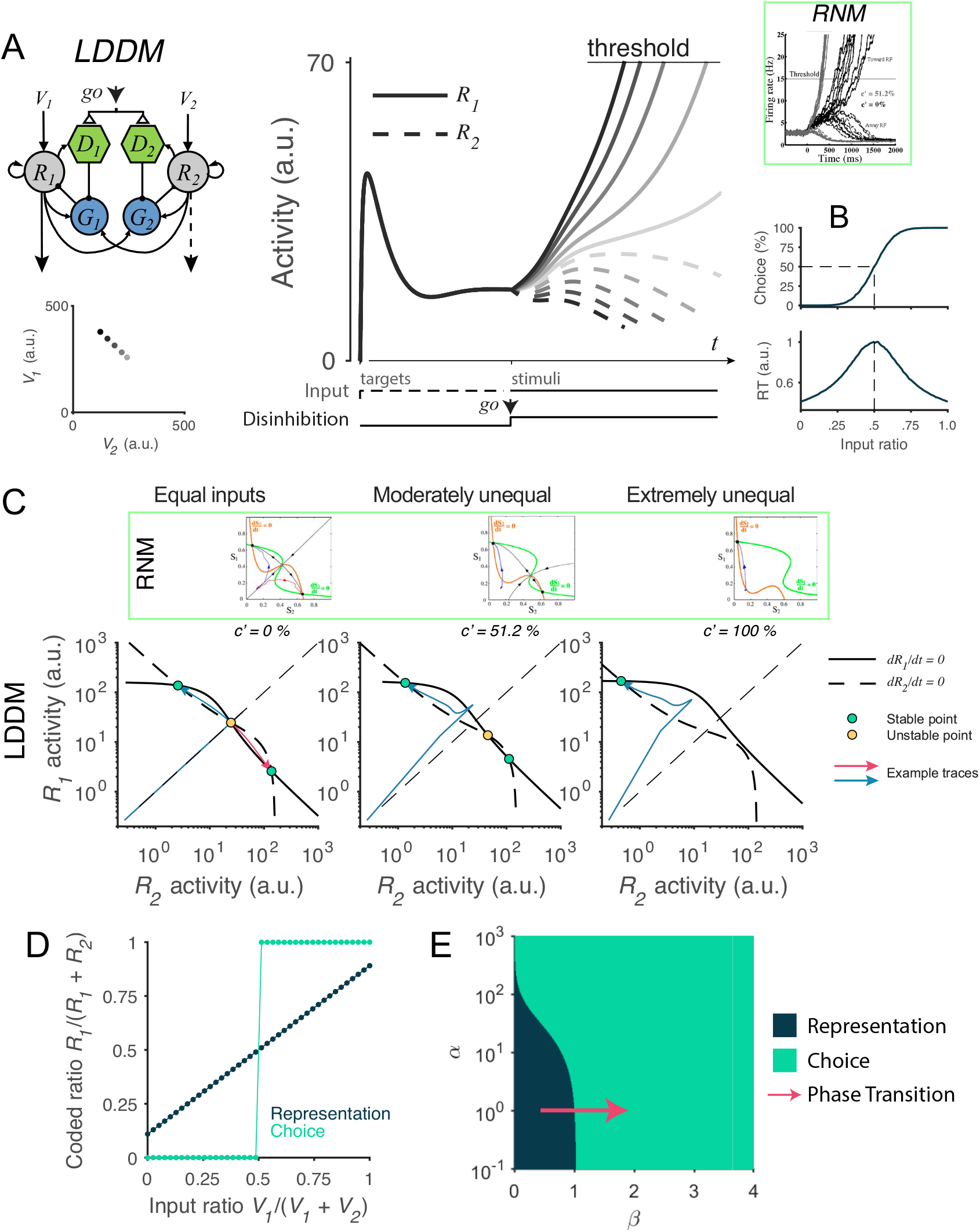
RNM-like WTA selection dynamics in the LDDM. **A**. Example *R*_*1*_ (solid) and *R*_*2*_ (dashed) dynamics in a classic reaction-time motion discrimination task. The model predicts phasic stimulus onset dynamics during the pre-stimulus stage and WTA dynamics during the stimulus stage when receiving different input values (left inset). Consistent with RNM dynamics (upper right inset), the *R* unit receiving stronger input ramps up to reach the decision threshold while the opponent *R* unit activity is suppressed; the speed of bifurcation depends on the input strength. **B**. Choice accuracy (top) and RT (bottom) predicted by the LDDM with noise. **C**. Phase-plane analysis of the LDDM (lower) compared with the original RNM (upper inset) shows basis for WTA dynamics under equal (left), moderately unequal (middle), and extremely unequal (right) inputs. Both models show similar features across input values: Under equal inputs, the nullclines of *R*_*1*_ and *R*_*2*_ intersect on three equilibrium points, with one unstable point (yellow) and two stable attractors (green) (left). Under unequal inputs, the basin of nullclines is biased to the side with stronger input (middle). When the inputs are strongly biased, only the attractor associated with stronger input retains (right). Red and blue lines show example traces of *R*_*1*_ and *R*_*2*_ activities. **D**. Comparison of the coded ratio between the representation (black) and WTA competition (green) regimes. While the LDDM preserves the input ratios during value representation, it shifts to a categorical coding of choice during WTA selection. **E**. Distinct normalized value coding (dark) and WTA competition (green) regimes in the parameter space defined by *α* and *β*. Acoss a wide range of *α*, the transition between valuation and selection regim1e4s can be implemented by an increase in *β* (pointed by red arrow). The insets in **A** and **C** were adapted from Wong & Wang (2006), Copyright 2006 Society for Neuroscience.

We find that the LDDM replicates neural and behavioral aspects of WTA competition. In **Fig. 5A**, we show example model activity for five input strengths corresponding to different motion coherence levels. Consistent with electrophysiological recordings in posterior parietal cortex^14,16,19^, model *R* unit activities bifurcate based on the input strengths, with the unit receiving stronger input ramping-up to an (arbitrary) decision threshold while the activity of the opponent unit is suppressed. The speed of bifurcation depends on the contrast between the inputs, a variable equivalent to motion coherence in the experimental literature^14,16^. When we simulated the LDDM with noise (additive Ornstein-Uhlenbeck noise; see Methods for details), the model reproduces the characteristic psychometric and chronometric functions that relate choice accuracy and reaction time (RT) to choice difficulty (**Fig. 5B**). Thus, the LDDM exhibits mutual competition that generates WTA selection, and reproduces empirical decision behavior previously described by RNM models (**Fig. 5A**, inset).

What features of the LDDM are essential to generate WTA competition? As shown in the phase-plane analysis (**Fig. 5C**), the network in the choice regime (*β*_0n_ = .9 in this example) shows a different configuration of nullcline intersections than the network in the value representation regime (*β*_off_ = 0; **Fig. 3B**). Given equal inputs, the nullclines of *R*_*1*_ and *R*_*2*_ intersect at three equilibrium points (left panel in **Fig. 5C**), with the central point unstable and the two peripheral points stable. Thus, given an initial configuration of *R*_*1*_-*R*_*2*_activities (in the presence of noise), the system will converge to the closer peripheral attractor (see example activity traces in blue and red thin lines) and implement WTA competition. Given moderately unequal inputs, the basin of attraction is biased towards the side with higher input, resulting in a higher probability falling into the side with higher input (middle panel in **Fig. 5C**). When inputs are extremely unequal, the stable equilibrium in the middle of the basin and the unstable equilibrium point associated with weaker input no longer exist, leaving only the attractor associated with stronger input (**Fig. 5C**, right). Thus, across varying degrees of input coherences, disinhibition drives the LDDM towards a selection of one of the potential choices. This can be seen in **Fig. 5D** by viewing the output ratio 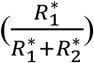of the preferred attractor as a function of input ratio 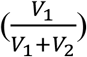: under active disinhibition we observe categorical coding (green line), in contrast to inactive disinhibition where the output ratio faithfully preserves the original ratio of inputs (dark line).

To understand the operating regimes of the LDDM, we quantified model behavior across the full parameter space defined by recurrent excitation weight (*α*) and local disinhibition weight (*β*), both of which are critical in determining the properties of the system (see *Equilibria and stability analysis of the LDDM* in **Supplementary Results** for the mathematical proof). Decisions with equivalent inputs are a critical test of WTA behavior, since WTA systems should select an option (stochastically) even in these symmetric scenarios^48–50,68^; we therefore analyzed system behavior under equal value inputs. As shown in **Fig. 5E**, this analysis revealed two distinct territories corresponding to value representation and WTA operating regimes. The value representation regime generates a unique attractor for normalized value representation but no WTA attractors; in contrast, the WTA regime (induced by a change in *β*) generates no normalization attractor but instead *R*_*1*_ and *R*_*2*_ always diverge into high-contrast attractors (see supplementary **Fig. S2** for a full description of regime parcellation). While the WTA regime asymptotically approaches zero disinhibition when recurrent excitation is extremely strong, local disinhibition is always required to generate WTA choice. Models with a wide range of recurrent excitation can transit from value representation to WTA choice with an increase in local disinhibition strength (for example, red arrow in **Fig. 5E**). These findings emphasize the importance of local disinhibition to WTA choice, and highlight a particular role for a dynamic gating signal in controlling the transition from value coding to option selection.

### The LDDM captures empirical choice behavior and neural activity

While the preceding analyses show that the LDDM can generate value normalization and WTA selection, a critical question is whether this circuit architecture accurately captures behavioral and neural aspects of empirical decision making. Here, we take advantage of the limited number of parameters in this differential equation-based LDDM (compared to more complicated conductance-based biophysical models^49,50,79,81^), which allows model fitting to empirical data. Specifically, we fit LDDM parameters to nonhuman primate behavior from the reaction-time version of the motion discrimination task described above. These choice and RT data from monkeys align with a reduced form model of decision making (the drift diffusion model)^120^, and the activity of posterior parietal neurons recorded during this task display characteristic decision-related features (motion-dependent ramping, a common decision threshold, and WTA activity).

To fit the LDDM to behaviorally observed RTs, we employed the standard quantile maximum likelihood method (QMLE) to the RT distributions across input coherence levels (0 – 51.2%), with correct and error trials dissociated^121–123^. After defining connection weights *ω*_ij_ as 1 for simplicity, the model is reduced to seven parameters: recurrent excitation weight *α*, local disinhibition weight *β*, noise parameter *σ*, input value scaling parameter *S*, and time constants *τ*_*R*_, *τ*_*G*_, and *τ*_*D*_ (see Methods for model-fitting details). Predictions of the best fitting model are shown in **Fig. 6A** (best fitting parameters: *α* = 0, *β* = 1.434, *σ* = 25.36, *S* = 3251, *τ*_*R*_ = .1853, *τ*_*G*_ = .2244, and *τ*_*D*_ = .3231). Model-predicted RT distributions (lines) closely follow the empirical distributions (bars) for both correct (blue) and error (red) trials across different levels of input coherence. The aggregated mean choice accuracy and RT data are shown in **Fig. 6C**. Model choice accuracy (line) captures the average empirical psychometric function (crosses); model RT captures coherence-dependent changes in the chronometric function, including longer RTs in error trials (dashed line and empty dots) compared to correct trials (solid line and dots). Beyond mean RT data, the LDDM accurately captured aspects of the empirical RT distributions, as evident in the quantile-quantile (Q-Q) plot of RT quantiles as functions of chosen ratio (**Fig. 6B**).

**Figure 6.**
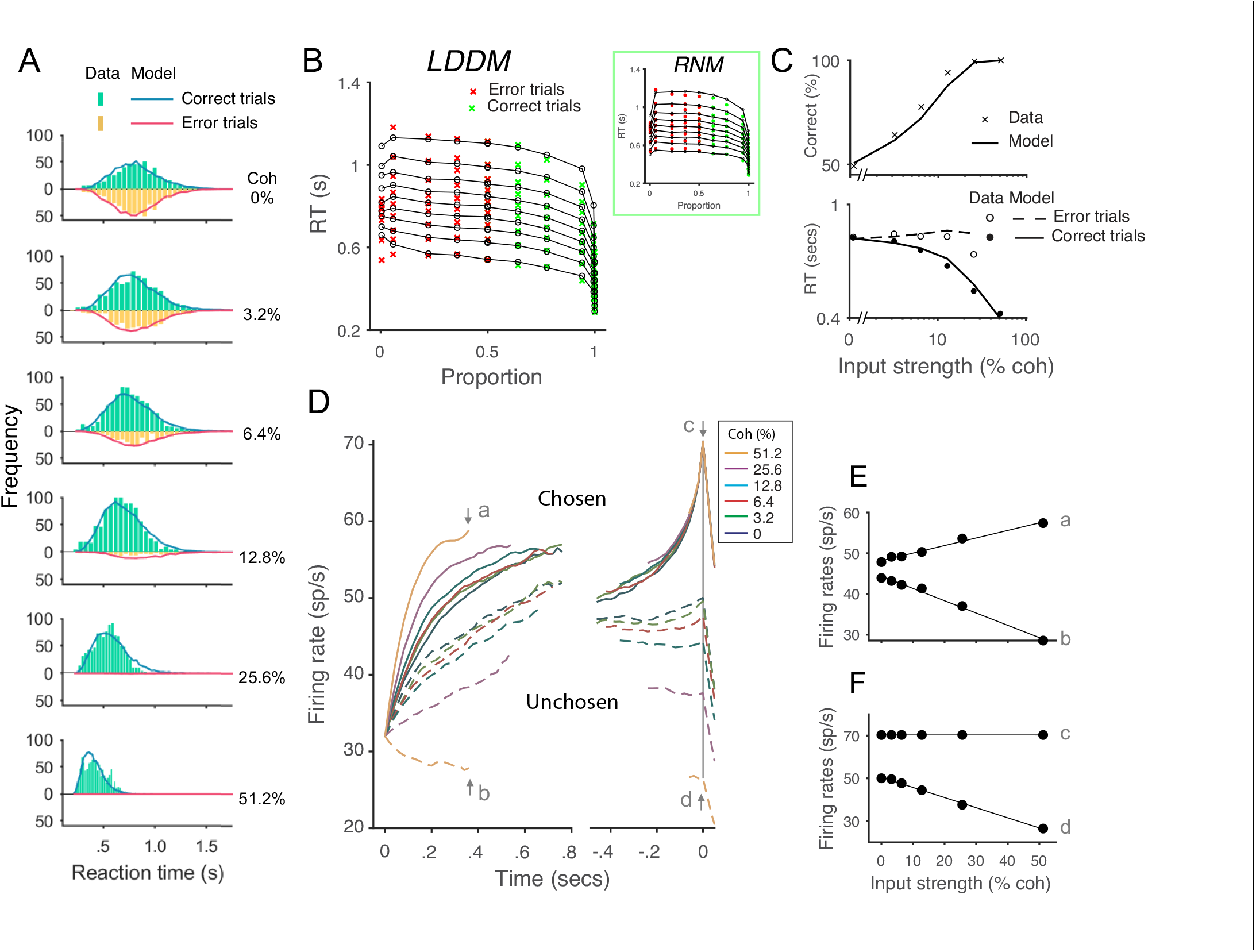
The LDDM performs as well as the RNM in capturing empirical behavior and neurophysiological data during perceptual decision-making. **A**. Model predicted RT distributions fit to behavioral data. Predicted RT distribution (lines) match the histogram of empirical RT distribution (bars), with correct and error trials separately (indicated by color) across levels of input strength (% coherence). **B**. The fitting results of the LDDM and the original RNM (upper-right inset) visualized in Q-Q plots. Nine quantiles of RT under each condition are stacked on the x-axis indicating the correct choice proportion under each input coherence (0 - .5 are error trials, shown in red cross; .5 - 1 are correct trials, shown in green cross). LDDM Model predicts the choice proportion and the shape of RT distribution as well as the original RNM. **C**. Model predicted psychometric function (upper) and chronometric function (lower). Choice accuracy aggregated by input strength (lines) fit well to the empirical data (crosses). The predicted RT aggregated by input strength for correct (solid line) and error (dashed line) trials capture well the RT for correct (filled dots) and error (empty dots) trials in empirical data. **D**. The best-fitting model replicates the neural dynamic features of the recorded neural activity. *R* unit activities aligned to the onset of stimulus inputs (left) and aligned to the time of model decision (right) replicate the stereotyped ramping dynamics of units associated with the chosen side (solid lines) and suppression of units associated with the unchosen side (dashed lines) under different levels of input strength. **E**. Effect of input strength on early stage model activity shortly after stimulus onset, at times indicated by arrows **a** (chosen side) and **b** (unchosen side). Consistent with empirical findings, the activity associated with chosen and unchosen options linearly increase and decrease with input strength. **F**. Effect of input strength on late stage model activity shortly before model choice, at times indicated by arrows **c** and **d**. Consistent with empirical findings, chosen unit activity reaches a common decision bound and show little input dependence, while unchosen activity remains suppressed by input strength. Empirical and behavioral and neural dataset from Roitman & Shadlen (2002).

We compared the performance of the LDDM in fitting this classical dataset with the reduced form of the RNM, which can be reduced to eight free parameters (see supplementary **Fig. S3** for details)^50^. The negative loglikelihood (nLL) and AIC values of the two models are close, with nLL_LDDM_= 16546, nLL_RNM_= 16573, AIC_LDDM_= 33109, AIC_RNM_ = 33165, suggesting that the LDDM performs as well as the original RNM in fitting behavior in the RT task.

Importantly, the LDDM – fit only to behavior – generates predictions about the underlying neural dynamics that can be compared to electrophysiological findings. We examined *R* unit activity in the best-fitting model, with predicted activity aggregated across trials and aligned to the onset of stimuli and the time of decision as in the original study^14^. Aligned to the onset of stimuli **(Fig. 6D**, left), neural responses are aggregated by coherence level and eventual choice, and truncated at median RT. These data show clear evidence of WTA competition: chosen (solid) and unchosen (dashed) activity traces diverge over time. Moreover, neural activity is stimulus-dependent: the dynamics of both chosen and unchosen units ramp at different, coherence-dependent speeds, consistent with empirical findings consistent with an accumulation process. More quantitatively, we examined the relationship between activity and coherence at the specific time point (arrow points **a** and **b**) reported in the original work (**Fig. 6E**). Model predictions align well with empirical observations: chosen activity shows a significant increase with input coherence (**a**, 18.56 spikes/second/100% coherence), while unchosen activity shows a decrease (**b**, -30.18 spikes/second/100% coherence).

Aligned to the onset of decision (**Fig. 6D**, right), model *R* unit activity near the time of choice shows further evidence of WTA competition observed in real neurons: the initial divergence between chosen and unchosen activity traces extends into a categorical coding of choice. The relationship between activity and coherence quantitatively replicates the empirical pattern immediately preceding the decision time^14^: chosen activity (indicated by arrow **c** in **Fig. 6D** and plotted in **Fig. 6F**) no longer shows much difference across coherence conditions (-.009 spikes /second/100% coherence), while unchosen activity (indicated by **d** in **Fig. 6D** and plotted in **Fig. 6F**) retains a decrease (**d**, -47.49 spikes/second/100% coherence). Thus, *R* unit activity – in a model with parameters fit only to behavior – replicates the recorded activity of parietal neurons during both initial decision processing and eventual choice selection.

### The LDDM integrates normalized value coding and WTA choices

While the LDDM separately replicates normalized value coding and WTA dynamics shown in different empirical studies, a key distinguishing feature of the LDDM is that it can capture both phenomena within a single experimental context. Numerous studies using the random-dot motion paradigm show two stages of dynamics: target (action) representation during the pre-motion stage and WTA selection after the go cue following motion stimuli^15,19^. Neural activity in the pre-motion stage shows a characteristic phasic-sustained dynamic to the presentation of visual cues; rather than purely sensory information, activity during this stage reflects the magnitude and probability of reward associated with the visual cues^15^. After the go cue, WTA dynamics reflects an integration of motion information and implements a transition from initial value coding to a categorical coding of choice in the late stage of decision^14–16,19,21,22^. Studies of economic choice show a similar set of dynamics, a context dependent valuation followed, after a go-cue, by a shift to WTA^2,12,17,27,29^. Interestingly, the number of alternatives affects the neural dynamics during both representation and choice^19,53,54^. When the choice set is expanded from two options to four options, early representational activity is lower during pre-motion dynamics (**Fig. 7A**) and the speed of WTA dynamics slows after motion onset (**Fig. 7C**).

**Figure 7.**
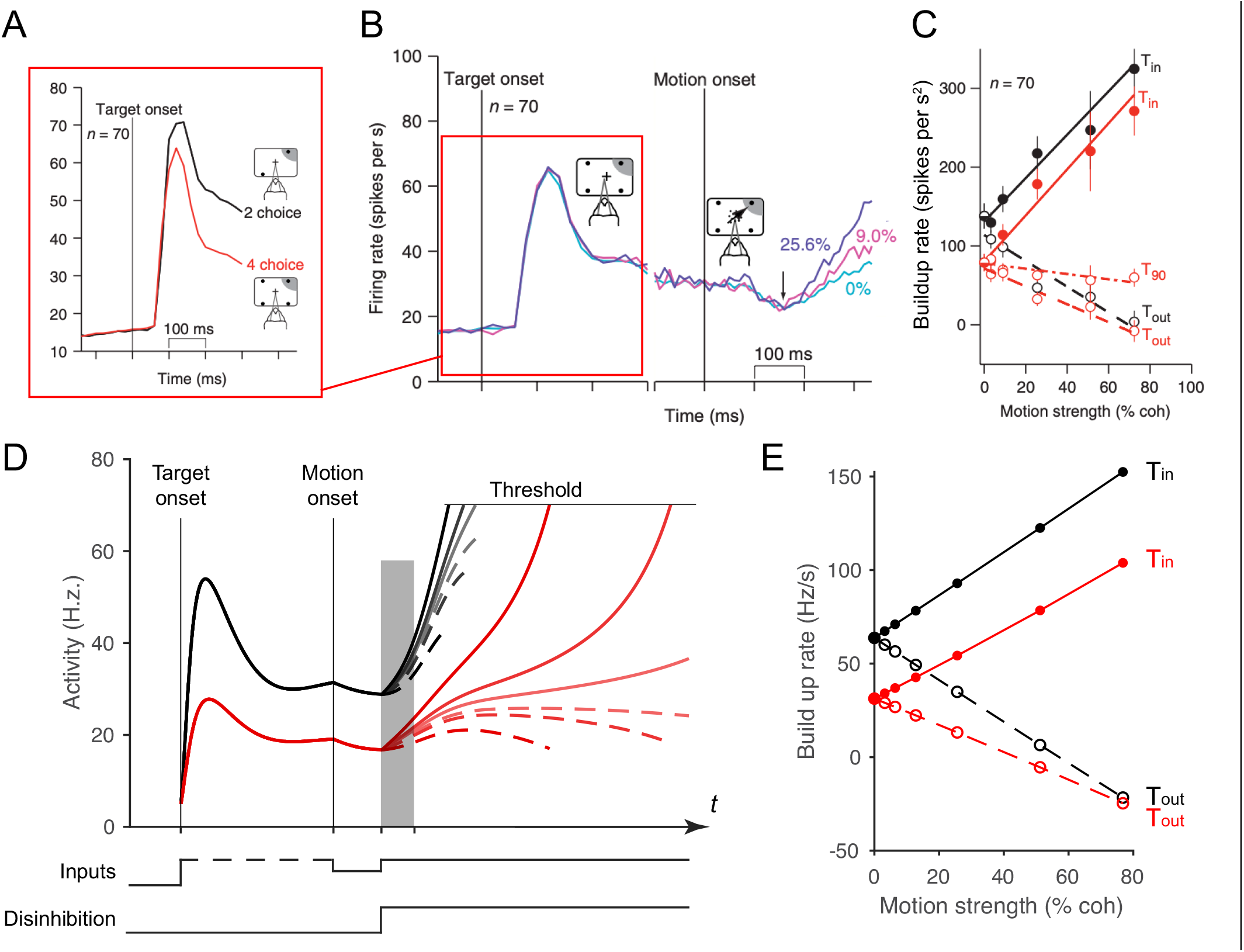
LDDM replicates both the normalized coding and WTA competition observed sequentially in single neurons examined in a multi-alternative choice task. **A**. Parietal neuron activity during pre-motion representation is decreased in 4-alternative (red) versus 2-alternative (black) trials. **B**. Neural activity during 2-alternative choice transitions from pre-motion target representation (left) and to post-motion onset WTA dynamics (right), shown for different input coherences (indicated by colors). **C**. Ramping speed in 2 (black) and 4 (red) alternative conditions, separated for choices towards (T_in_) and away from (T_out_) the neural response field (T_90_ in the original study designates choices for targets orthogonal to the Tin-Tout target, and is not examined here). **D**. Dynamics of LDDM *R* unit activity during pre-motion representation without disinhibition (left) and after motion onset with disinhibition (right). **E**. LDDM replicates the decrease in ramping rates (time period shaded in **D**) from 2 (black) to 4 (red) alternatives after the motion onset, consistent with the empirical data. Panels **A – C** are adapted with permission from Churchland et al., 2008.

Accordingly, in this section we examined whether the LDDM replicates the impact of the number of alternatives on both early and late empirical neural dynamics during both the representation phases and the WTA phases observed in real neurons. Under four (versus two) alternatives, LDDM R unit activity during the representation stage decreases because of increased recurrent inhibition, driven by multiple contextual inputs (left side in **Fig. 7D**). Similarly, the ramping speed after motion onset and disinhibition decreases in the 4-alternative (versus the 2-alternative) condition, despite identical parameters (**Fig. 7E**). These results highlight the LDDM as a potential mechanism of integrating normalized value coding and WTA competition within a single circuit architecture.

### A novel form of persistent activity

We next examine implications of the local disinhibition architecture for another characteristic of decision-related neural firing: persistent activity. In cortical areas such as parietal^10,14,16,21,124^, prefrontal^125–128^, and premotor cortices^2^, neurons show elevated firing in the absence of stimulus-driven input over intervals of seconds; such persistent activity is thought to underlie working memory and enable decisions based on internally maintained information. In RNMs, recurrent excitation and feedback inhibition preserve categorical choice information after input withdrawal because of point-attractor dynamics^49,50,68^. Here, we answer two questions: does the LDDM generate persistent activity, and how does this persistent activity differ from that in standard RNMs?

We found that the LDDM can generate two distinct forms of persistent activity, controlled by the state of disinhibition. **Fig. 8A** shows example dynamics of two *R* units before and after withdrawal of inputs while disinhibition is silent. Following input withdrawal, network activity decreases but still preserves elevated firing rates, governed by the self-excitation parameter *α* (the network loses elevated activity when *α* ≤ 1). The persistent activity ratio between *R*_*1*_ and *R*_*2*_preserves the ratio between the input values *V*_*1*_ and *V*_*2*_, in contrast to RNMs which only preserve categorical information about the largest value (see supplementary **Fig. S5** for proof). Phase-plane analysis suggests that relative value coding in persistent activity arises from a line-attractor dynamic in the network during the inactivation of disinhibition, unlike point-attractor dynamics in the RNM (**Fig. 8B**). Like other line-attractor models of persistent activity that store continuous-valued information^74,129–131^, an unbiased coding of the input ratio requires perfectly balanced gain control weights from *G* to *R*. Unbalanced weights will result in distorted coding of the input ratio and graded coding of the inputs will decay over time (supplementary **Fig. S5**). For perfectly balanced weights, the line attractor state is vulnerable to noise perturbation. A small perturbation can easily drive the activity to drift on the line of attractors, with the summed value of *R*_*1*_ and *R*_*2*_as a constant 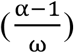 The preserved ratio between *R*_*1*_ and *R*_*2*_ drifts stochastically over time, similar to the prediction of other line-attractor circuits and consistent with behavioral and neural variability related to working memory^130,132^.

**Figure 8.**
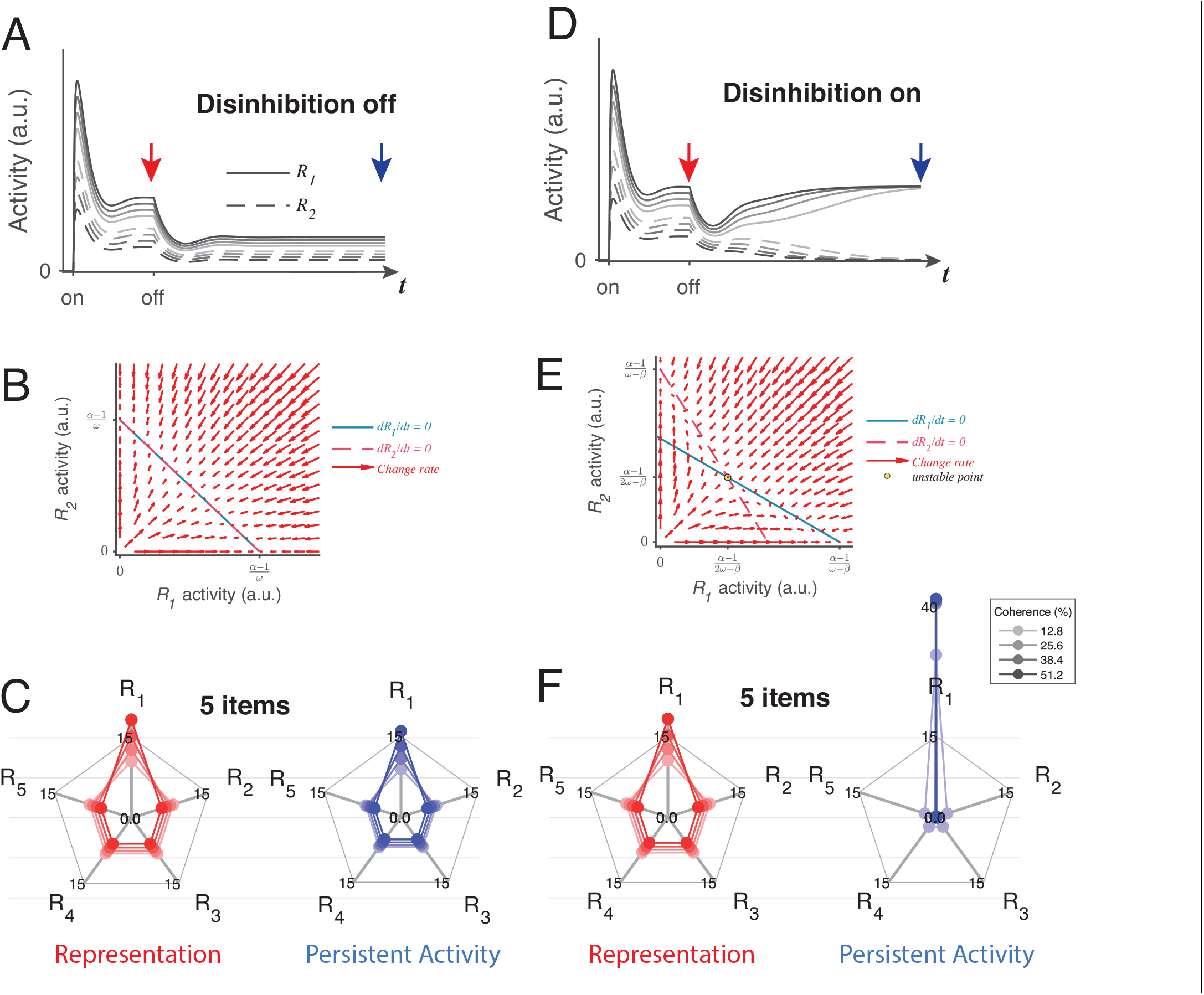
LDDM disinhibition controls flexible implementation of either line attractor or point attractor dynamics in persistent activity. **A-C**. LDDM under silent disinhibition preserves the input ratio information during persistent activity. **A**. Example *R*_*1*_ (solid) and *R*_*2*_ (dashed) activities before and after withdrawal of stimuli under different levels of inputs. Neural activity decreases after withdrawal but reaches a new steady that preserves graded coding of the inputs. **B**. Phase-plane analysis of persistent activity exhibits a line attractor under inactivated disinhibition. The nullclines of *R*_*1*_ (blue) and *R*_*2*_ (red) intersect on the line of attractors, on which the summed value of *R* activities is a constant 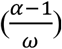. Red arrows indicate the instantaneous change rate of *R*_*1*_ and *R*_*2*_ at given initial values, following the direction that preserves the *R*_*1*_-*R*_*2*_ ratio. **D-F**. Persistent activity under active disinhibition preserves only them largest item as categorical information. **D**. Example *R*_*1*_ and *R*_*2*_activities before and after withdrawal of stimuli. Disinhibition activates at the same time as the offset of stimuli. During the delay period, the activity dynamic gradually switches from a graded coding of the inputs to a WTA type of categorical coding, preserves only the larger item. **E**. Delay period phase-plane analysis exhibits a point-attractor state under activated disinhibition. The nullclines of *R*_*1*_(blue) and *R*_*2*_ (red) intersect on an unstable point. Red arrows indicating the instantaneous change rate of *R*_*1*_ and *R*_*2*_ bifurcate from the middle to the side corners, resulting in a high-contrast categorical coding. **C** and **F**. Expansion of the LDDM from a 2-item circuit to a 5-item circuit, under inactivation and activation of disinhibition. Each axis on the radar plot indicates the activity of one *R* unit. Dots connected with a line indicate the *R* activities under the same input conditions. The input values change according to coherence level (c’) as *S*\*[1+c’] for *R*_*1*_ and *S*\* [1-c’] for *R*_*2*_-*R*_*5*_. Representation before withdrawal of inputs (left panels in **C** and **F**) and persistent activity without disinhibition (right panel in **C**) preserve the information about the input values. While persistent activity under disinhibition (right panel in **F**) only preserves the item received the largest input, with activities of the other items suppressed.

However, a line attractor is not the only state that the LDDM predicts. If disinhibition is activated during the delay interval, the network switches to a point attractor dynamic (see supplementary **Fig. S6** for proof). **Fig. 8D** shows example dynamics of two *R* units before and after withdrawal of inputs. Disinhibition drives a competition between the two *R* units, resulting in a switch between graded coding of the input ratio to a categorical coding of the largest value (*β* = .4 in visualization). Interestingly, a transition of coded information from input values to categorical information has been widely observed in firing rates in decision related regions, such as LIP and superior colliculus, during the delay period of decision making^15,16,55^. The point attractor predicted by the circuit under disinhibition (**Fig. 8E**) is highly tolerable to perturbations compared to the line attractor, and choice performance over long delays may require a switch from the value coding to the categorical regimes. As a plausible biological mechanism for mediating top-down control, disinhibition may gate such a transition without changing the network architecture.

The LDDM can be easily expanded to multiple alternatives. Here we show an example of a 5-alternative case, with 5 sets of option-specific *R*-*G*-*D* units. A line attractor network with silent disinhibition (**Fig. 8C**, right) is able to retain input value information of the 5 items simultaneously in the network. Due to normalization, the neural activity representing each alternative decreases with the total number of alternatives, with the summed value as a constant 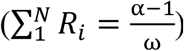, leading to a lower signal-to-noise ratio when coding more items; this set-size effect may be related to WM memory span constraints^133–137^. When disinhibition is active, the LDDM exhibits a point attractor (**Fig. 8F**, right), and the network only holds the information of the largest item as a categorical code during persistent activity.

### Gated disinhibition provides top-down control of choice dynamics

In addition to its crucial role in generating WTA competition, local disinhibition provides an intrinsic mechanism for top-down control of choice dynamics. Decision circuits show remarkable flexibility in timing, with similar neurophysiological data recorded in a variety of task paradigms: in addition to reaction-time tasks, in which subjects can choose at any time immediately after onset of stimulus, decision-related neural activity has been widely studied in fixed-duration and delayed-response tasks. In fixed-duration tasks, subjects are required to withhold selection of an action until an instruction signal. Neural activity prior to the instruction signal reflects value information, for example about reward characteristics^13,17,18,27,138^ or accumulating perceptual evidence^3,10,14–16,21,124^; however, this activity never fully diverges or reaches the decision threshold until after the instruction cue, suggesting a gating of the selection process. In delayed-response (working memory) tasks, subjects must postpone selection for an interval that includes both stimulus presentation and an additional subsequent interval after the stimulus is withdrawn. As in fixed-duration tasks, neural activity in delayed-response tasks typically carries decision–related information (across both the stimulus and delay periods) but WTA selection – and behavioral choice – is withheld until the instruction cue is given^3,10,14,16,21,124^. Thus, biological decision circuits are able to evaluate choice options while selectively initiating the WTA selection process with variable context-dependent timing.

How neural circuits implement dynamic control of selection – and a temporal separation of evaluation and WTA choice – is largely unaddressed in current decision models. In RNM models, neural activity is driven by attractor dynamics; option evaluation and the selection process cannot be disambiguated, and WTA competition is not under top-down control. Here, we examine how the timing of a dynamic top-down control signal – modulating the strength of disinhibition via long-range inputs and neuromodulation – allows the LDDM to capture neural activity in different task paradigms. In these simulations, disinhibition is activated when the instruction cue to choose is presented. **Fig. 9A** shows LDDM activity in a reaction-time task, a standard paradigm in perceptual decision-making^14,19^. As in prior analyses (**Figs. 5** and **6**), LDDM *R* units show simultaneous evaluation (coherence-dependent ramping) and WTA selection (rise to threshold) processes, driven by an immediate activation of disinhibition at motion stimulus onset.

**Figure 9.**
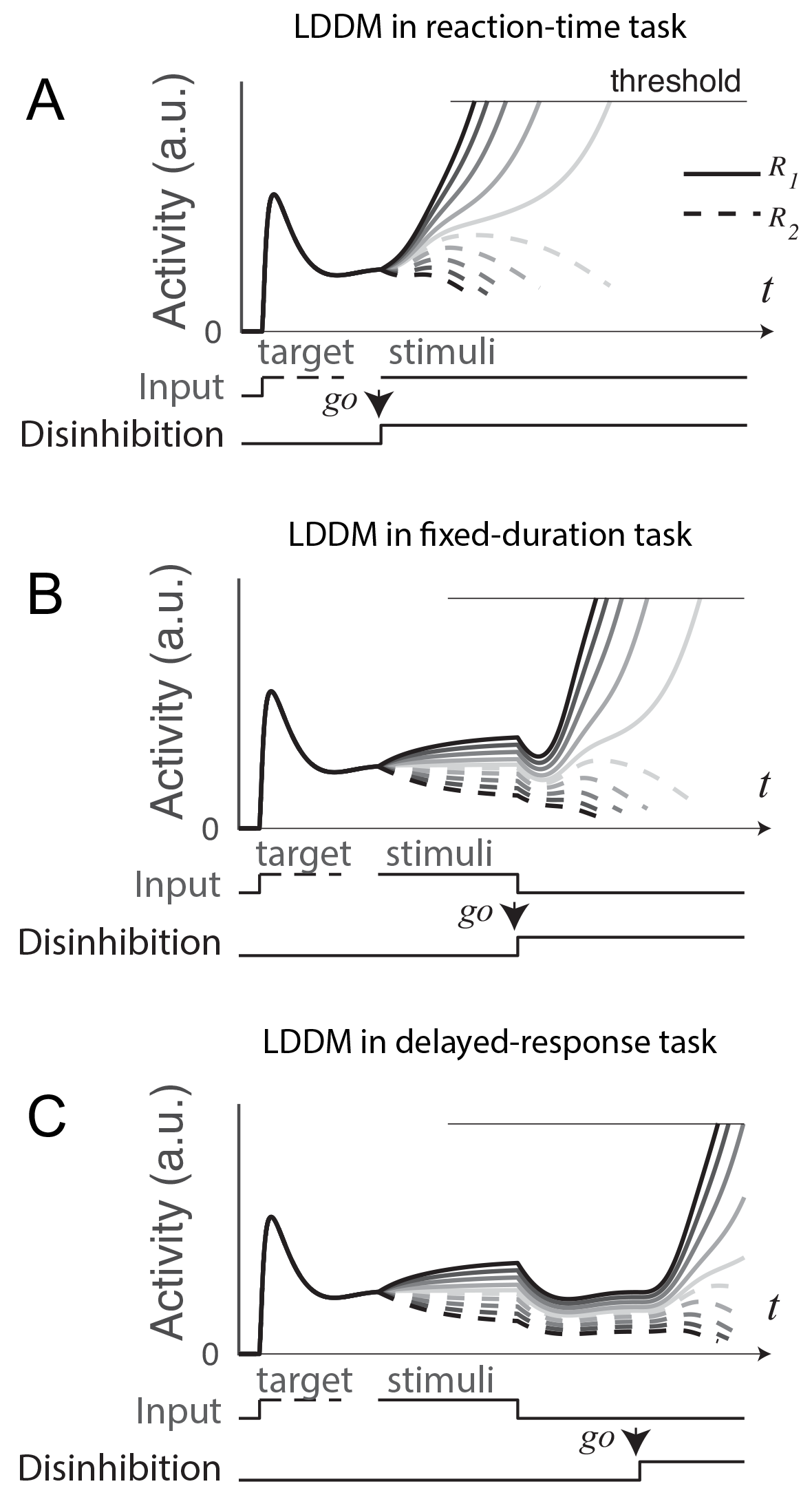
Gated disinhibition flexibly adapts the dynamics of the circuit to various types of tasks. All of the tasks consist of a pre-stimulus stage with equal inputs to *R*_*1*_ (solid) and *R*_*2*_ (dashed) and a stimulus stage with input values determined by the stimuli (indicated by grayscale, the same value matrix as used in Figure 5A). **A**. Reaction-time task. Subjects are free to respond at any time following stimulus onset, and model disinhibition is activated with the onset of stimuli. Model dynamics show WTA competition right after the onset of stimuli. **B**. Fixed duration task. Subjects are required to wait for a fixed duration of stimulus viewing before choice, and model disinhibition is turned on only at the onset of the instruction cue (usually indicated in experiments by fixation point offset). Model dynamics show normalized value coding before the instruction cue and a transition to WTA choice afterward. **C**. Working memory (delayed response) task. Subject choice occurs after an interval of stimulus presentation and a subsequent delay interval without stimuli, and model disinhibition is turned on at the end of the delay period. Model dynamics exhibit normalized value coding during stimulus input, preserved relative value information during the delay period, and a transition to WTA choice dynamic after the instruction cue.

In a fixed-duration task (**Fig. 9B**), disinhibition is activated after a required interval of stimulus presentation. Compared to the reaction-time task, LDDM activity here shows distinct, temporally separated patterns during stimuli viewing and option selection; this temporal segregation is driven by the activation of disinhibition (a step function on *β* in this example), which promotes a transition between value representation and WTA choice.

A further demonstration of this temporal flexibility arises from considering delayed-response tasks (**Fig. 9C**), which include an interval between stimuli offset and onset of the instruction cue. Consistent with its ability to maintain persistent activity (**Fig. 8**), the LDDM shows value coding across the delay interval and implements WTA selection until instruction and accompanying activation of disinhibition. These results show that the LDDM – via modulation in the timing of disinhibition activation - can temporally separate the value representation and selection processes, enabling it to capture the diversity of neural dynamics seen in reaction-time, fixed-duration, and delayed-response tasks.

### GABAergic potentiation distinguishes LDDM from earlier models

The architecture of disinhibition employed by the LDDM is more structured than earlier non-selective inhibition used in more standard competition networks. This distinction gives rise to the novel prediction from LDDM that the influence of global changes in inhibitory tone are non-selective during representation, but switch to being input-selective after disinhibition is increased. This reflects a fundamentally novel prediction of this class of model. To empirically test that key prediction, optogenetic/pharmacological manipulation of GABAergic activity could be introduced. The LDDM contains two different types of inhibition and thus its reaction to a GABAergic agonist depends on both the state of the disinhibitory network and the intensity of the GABAergic activation. To highlight the importance of that prediction, we implemented different levels of GABAergic activity in both the LDDM and the more traditional RNM.

At the neural level, the LDDM predicts a dissociable effect of increased inhibition on excitatory neuron activity (**Fig 10A**). During option representation (cue interval in fixed duration trials), increased inhibition increases both recurrent and lateral inhibition, leading to decreased excitatory neuron firing rates and a weaker modulation by value. During option selection (go/choice intervals in fixed duration trials), stimulation of local disinhibition increases WTA activity but decreases the late-stage representation of value. At the behavioral level, these changes should produce a speeding up of RTs and a decrease in choice accuracy (**Fig. 10B**), expected differences between control and agonist that would be evident in chronometric and psychometric curves (**Fig. 10C**). Note that the qualitative predictions for agonist effects on RT and accuracy (i.e. direction of change) are robust to specific LDDM parameterizations (**Fig. 10D**). In contrast, in more traditional networks like the RNM that employ non-selective inhibition, increased inhibition suppresses both the baseline and WTA stages of neural activities (**Fig. 10E**). The suppression in neural coding will slow down RTs but will not affect the choice accuracy (**Figs. 10F** and **G**). These novel predictions could be readily tested and differentiate models that rely on the structured disinhibition that we propose from models that employ more traditional changes in the E/I balance to achieve state changes.

**Figure 10.**
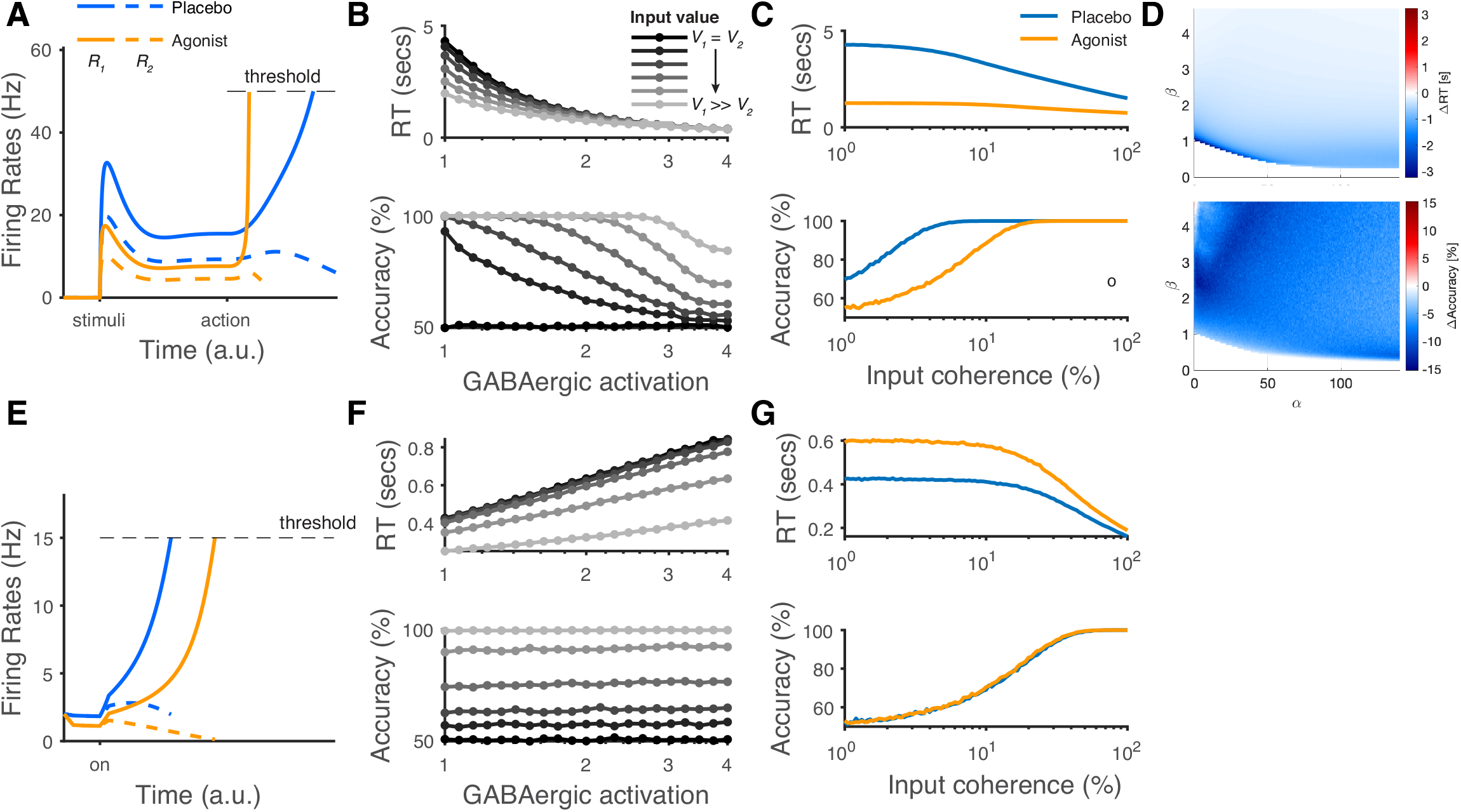
The modelling predictions of GABAergic agonist to decision-making neural dynamics and behaviors. **A**. The predicted neural dynamics of pyramidal neurons (R1, solid lines and R2, dashed lines) activities in a fixed duration decision task from LDDM. Agonist (orange) comparative to control (blue) decreases neural activities during early stage representation but speeds up WTA bifurcation after during choice. **B**. GABAergic activation speeds up RTs but decreases choice accuracy, examined over multiple levels of input coherences (indicated by gray scales). **C**. Comparing GABAergic agonist condition (orange) with control (blue), the differences should be evident in average chronometric and psychometric curves. **D**. The predicted behavioral pattern can be generalized across the full space of *α* and *β* parameters regime. **E**. The predicted neural dynamics of pyramidal neurons (R1, solid lines and R2, dashed lines) activities from RNMs (e.g., Wong and Wang, 2006). Since the model does not include a mechanism of switch, fixed duration task is not able to be tested in this type of model. We examined the reaction time task instead. RNM predicts suppressed neural dynamics under increased GABAergic activity. **F**. RNM predicts increased RTs but un changed accuracy. **G**. The chronometric and psychometric curves predicted by RNM will be qualitatively different from LDDM.

## Discussion

Normalized value coding, WTA choice, and persistent activity are three important and characteristic features of decision making which have all been observed in single neurons. We find that a hybrid model that unifies these key features can be implemented with a specific biologically plausible architecture – local disinhibition. In the LDDM, gated disinhibition separates the value normalization and WTA choice computations, enabling them to be generated in the same circuit architecture. Notably, the gated disinhibition in the LDDM replicates features of diverse, existing computational models: the top-down control of normalization via disinhibition mirrors recently proposed mechanisms for flexible modulation of contextual processing in sensory circuits^139–141^ and input-scaled disinhibition implements a self-sparing inhibition motif central to midbrain models of categorical selection^142,143^. When fit to empirical behavioral observation, the LDDM accurately captures choice and RT patterns, driven by underlying model dynamics that reproduce the neural dynamics of empirical neurophysiological findings. Since the vast majority of empirical neural responses have been recorded from putative pyramidal neurons, we focus here on excitatory LDDM responses; however, the structured inhibition we model from newer anatomical data predicts input-selective inhibition. The model also makes novel predictions about inhibitory and disinhibitory activity dynamics and pharmacological manipulations that may warrant future examination. Furthermore, via disinhibitory control, the LDDM can exhibit both line attractor and point attractor forms of persistent activity without a change in the circuit structure. Finally, gated disinhibition in the LDDM provides a mechanism for top-down control of decision dynamics. Controlling the timing of disinhibition paces the decision process and replicates neural dynamics from various choice task variants.

While normalized value coding and WTA selection have largely been modeled separately, the LDDM offers a biologically-plausible circuit architecture that integrates the two features. Existing neurophysiological evidence show that WTA dynamics and normalized coding co-exist in the same brain regions. On the one hand, neural activities show relative value coding in the early stage of decision-making, reflecting a context-dependent modulation consistent with the canonical divisive normalization computation^2,9,15,19,26,27,30^. On the other hand, WTA choice dynamics are widely observed during decision making across multiple brain regions of non-human primates^2–25^, including many of the brain regions that show normalized value coding. In addition, a transition from graded coding to WTA choice has been widely documented in the decision relevant regions mentioned above. Neural firing rates shows a graded coding of perceptual evidence and reward during the early stage of decision-making and gradually transition to a categorical coding for choice in the late period of decision-making^13–19,45,55,144^. However, the evidence for one alternative is typically inversely related to the evidence for the other alternative, making it difficult to dissociate the dynamic effects of evidence integration and contextual information about other alternatives.

In the LDDM, disinhibition modulates the dynamics of the circuit without requiring changes in circuit structure. Existing models capture activity dynamics only in specific temporal intervals during decision-making tasks, or across trials in specific task paradigms^29,31,49,50,76^, and thus typically do not generalize across tasks. In contrast, gated disinhibition in the LDDM – driven by the external action instruction cue - controls the timing of valuation-to-WTA regime transition, enabling the LDDM to replicate neural dynamics in diverse task paradigms with different stimulus and action timing schedules^14–16,21^. Recent research on neuromodulatory control of disinhibition offers biologically plausible mechanisms for such top-down control of circuit dynamics. In addition to evidence that VIP neurons are recruited by long-range projections from distanced regions^100,114^, VIP neurons are recruited by neuromodulatory projections such as acetylcholine^105^ from the basal forebrain and pedunculopontine nuclei and serotonin from the red nucleus. With ionotropic acetylcholine receptor (nAChR) and serotonin receptors (5HT_3a_R and 5HT_2_R), VIP neurons depolarize to acetylcholine and serotonin^96,97,102,111^. The spiking mode of a major type of VIP neurons in layer II/III of the cortex switches from an input-insensitive burst-quiescent mode to an input-sensitive tonic mode under cholinergic and serotonin modulation^112^. Such a mode-switching feature allows the disinhibitory neurons to receive excitatory projections with different gain under different level of neuromodulation, providing a mechanism to modulate network dynamics via disinhibition without a change in network structure. *In vivo* studies show that disinhibition mediated by cholinergic activation is triggered in a surprisingly fast time scale of tens of milliseconds^101,111^, supporting a fast modulation mechanism of disinhibition and network plasticity.

An interesting feature of the LDDM is that it can produce both point attractor^145–148^ and continuous/line attractor^129,132,149^ dynamics in persistent activity, a balance controlled by the level of disinhibition. Given ambiguous empirical evidence, it remains controversial whether persistent activity in neural circuits exhibit point attractor^145–148^ or continuous/line attractor^129,132,149^ dynamics, and existing circuit models of persistent activity exclusively predict either a point attractor^73,81,150,151^ or line attractor^74,129–131,152^. The LDDM can generate both line attractor and point attractor states, suggesting that attractor dynamics might not be a fixed property of a network; rather, it may be adaptive and controllable by a top-down signal operating via gated disinhibition.

In conclusion, here we introduce a novel, biologically-plausible architecture for decision making based on local disinhibition, unifying the characteristic decision-making features of normalized value coding, WTA competition, and persistent activity into a single circuit. The LDDM captures psychometric and chronometric aspects of behavioral choice and predicts realistic neural dynamics in standard decision-making tasks. Local disinhibition provides a mechanism for top-down control of local decision circuit dynamics, enabling the LDDM to replicate variable task-dependent timing in diverse decision-making paradigms and implement speed-accuracy tradeoffs. These results suggest a new circuit mechanism for decision making, and emphasize the importance of incorporating interneuron diversity, local circuit architecture, and top-down control into models of the decision process.

## Methods

### Numerical simulations

To quantify neural dynamics and behavioral performance (choice/RT), time varying activity was represented by a system of differential equations (see Results) which was solved numerically using the Runge-Kutta method implemented in MATLAB (MathWorks) at time step of 1 ms. Evaluations using smaller time steps (0.1 ms) were examined and produced similar results. At each time step, the model unit activities were updated based on their values at the previous step according to the differential equations. Considering the biological reality that spike rates cannot be negative, the activities were constrained to be non-negative. For the simulations including noise, we assumed an additive noise term for each unit, which evolved independently based on an Ornstein-Uhlenbeck process (Eq. 5),

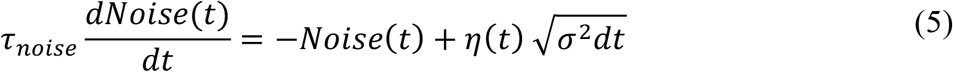

where *σ*^2^is the variance of the noise, *η* is a Gaussian white noise with zero mean and unit variance, and *r*_*noise*_ is the time constant for the noise fluctuation process. The time constant for the noise process (*r*_*noise*_) was set to 2 ms aligned with previous studies^49,50^. Note that this approach assumes for convenience that noise arises in model unit activity; however, similar stochasticity can be implemented assuming noise arises in inputs external to the circuit, generalizing our findings.

All parameters used for visualization were set as following unless specified elsewhere or fitted as free parameters in **Fig. 6**: *τ* _*R*_, *τ* _*G*_, and *τ* _*D*_ were set as the same value of 100 ms; the gain control weight *ω*_%3_ was set as unit value of 1 for simplicity; the self-excitation weight *α* was set as 15; the disinhibition weight *β* was assumed as zero in representation (i.e., *β*_off_ = 0) and set as 1.1 in WTA competition; the input values *V*_*1*_and *V*_*2*_were set as *S*^*^(1+c’) and *S*^*^(1-c’), where c’ indicates the motion coherence of the stimulus, with varied values [0, 3.2, 6.4, 12.8, 25.6, 38.4, 51.2]%) and *S* indicates the scale of input (set as 250). Baseline input *B* was set as 70 in **Figs. 3** and **5** and set as 0 in **Figs. 7-9**. Ornstein-Uhlenbeck noise was set as zero in most of the simulation but σ = 10 in **Fig. 5B**. In **Fig. 7**, some parameters are adjusted by definition (but not fit) to predict the multi-alternative choice data. *α* was set as 0; *β* was set as 1.5; scaling parameter was set as 640 for both pre-motion and motion period but set as 427 for the first 190 ms of motion period to replicate the initial dip. *τ* _*R*_, *τ* _*G*_, and *τ* _*D*_ were set as 100 ms. All parameters were kept the same between 2 and 4-alternative choices. In **Fig. 8**, parameters are adjusted between 2-item and 5-item cases in order to get comparable scale of activities in visualization: for 2-item case, *S* = 250, *α* = 15, and *β*_0n_ = .4; for 5-item case, *S* = 50, *α* = 37.5, and *β*_0n_ = .1.

### Fit the LDDM to empirical behavioral data

The LDDM with seven free parameters (the weights of self-excitation (*α*) and disinhibition (*β*), the variance of Gaussian white noise in the Ornstein-Uhlenbeck process (*σ*^2^), the scaling parameter of input (*S*), and time constants for three types of units *τ* _*R*_, *τ* _*G*_, and *τ* _*D*_) was fit to choice behavior (RT and choice accuracy) in a classic perceptual decision-making dataset^14^. We employed the commonly used quantile maximum likelihood estimation (QMLE) method^122,153^. The rationale of QMLE is to minimize the differences between the predicted data and the empirical data on the proportion of number of trials located in each RT bin. Choice accuracy was implicitly estimated because the algorithm accounts for the proportion of number of trials between correct and error trials. Nine quantiles (from .1 to .9 with .1 of step size) were used resulting in ten RT bins, with correct and error trials accounted for separately at each coherence level. Because the LDDM has no closed-form analytic expression for the RT distribution, we evaluated the prediction by Monte Carlo simulations (10240 repetitions for each input coherence). In each simulated trial, the initial values of *R*_*1*_ and *R*_*2*_ activities were set as 32 Hz to be comparable to empirical data^14,19^. Visual stimulus (motion) inputs were defined as *S*^*^ (1+c’) and *S*^*^(1-c’) for *V*_*1*_ and *V*_*2*_, where the free parameter *S* models input scaling and the coherence c’ replicated values in the original experiment (0, 3.2, 6.4, 12.8, 25.6, and 51.2 %)^14^. At visual stimulus onset, a gap period (90 ms) was implemented in order to capture the commonly observed initial dip in empirical firing rates^14^. Gated disinhibition was activated along with inputs after the gap. A decision was reached when either of the *R* unit activities reached a decision threshold of 70 Hz, the biological threshold observed in the empirical data^14^. 30 ms was added to the RT of threshold hitting to capture the delay in the down-streaming motor execution. After the decision, the input values, self-excitation, and disinhibition were reset to zero. The negative loglikelihood (nLL) of QMLE was minimized using Bayesian adaptive direct search algorithm in Matlab^154^. The estimation was conducted using GPU (NVIDIA Tesla V100) parallel computation on a high-performance cluster (NYU Langone), with 160 chains of random initial parameter values to prevent local minima. The chain with the smallest nLL in its fitting result was selected as the best fitting result.

The visualization of the predicted RT distribution (**Fig. 6A**) was calculated based on 60 evenly distributed RT bins, with correct and error trials calculated separately under each coherence. The predicted neural dynamics (**Fig. 6D**) were generated using the model best fit to behavior. *R* unit activities were aggregated across correct trials, segregated by units associated with the chosen side and unchosen side. As in the original experiment data visualization^14^, activity early in trials was aligned to stimulus onset and data within 100 ms of boundary crossing were omitted to reduce the impact of decision dynamics on visualizing early stage ramping dynamics. Early activity traces were cut off at the median value of RT for each coherence level to ensure that the average trace was based on at least half of the trials. Activity late in trials was aligned to the time of decision, and data within 200 ms of stimulus onset was omitted.

### Fit the LDDM and the DNM to the neural firing rates of normalized value coding

In order to quantify the performance of the LDDM in fitting to the neural dynamics of normalized value coding and compare with the original DNM, we fit the equilibrium values of the LDDM and DNM to the dataset of normalized value coding (**Fig. 4** in Louie et al., 2011). In this task, monkeys are asked to represent the reward targets (1, 2, or 3) on the corresponding location of the screen. The neural activity in the response field receiving direct input *V*_*1*_ is recorded. Different combinations of *V*_*1*_, *V*_*2*_, and *V*_*3*_ are provided to the monkeys based on the associated volumes of juice in the presented targets (varying from 50, 100, 200, and 250 µl or omitted target marked as 0; see details in Louie et al., 2011), resulting in 28 data points. The LDDM and DNM are expanded to a trinary-choice circuit, with *R*_*1*_, *R*_*2*_, and *R*_*3*_ receiving input from *V*_*1*_, *V*_*2*_, and *V*_*3*_, respectively. One free parameter is fitted for the DNM (Setting *α* = 0 reduces the form of the LDDM in Eq. 1 to DNM; Baseline input *B* in Eq. 1 is fitted). And two free parameters are fitted for LDDM (*α* and *B* in Eq. 1). The fitted neural firing rates were further rescaled by a scaling parameter *S* to match the rescaled empirical data points (**Fig. 4B**). Other parameters in Eq. 1 – 3 are set as following: The dimensionless values of *V*_*1*_, *V*_*2*_, and *V*_*3*_ set equals to the volumes of juice reported in the original paper, *β* = 0, *ω*_*ij*_ = 1, *τ* _*R*_ = *τ* _*G*_ = *τ* _*D*_ = 100 m*s*, and Ornstein-Uhlenbeck noise *σ* = 0. Bayesian adaptive direct search algorithm (BADS) was implemented to minimize the ordinary squared error between the steady state of the predicted neural firing rates on *R*_*1*_ and the empirical data.

### Simulation on pharmacological manipulation

In **Fig. 10** we tested GABAergic agonist manipulation effects in both the LDDM and RNM ^50^ by assuming different levels of enhancement of the GABAergic projections from the inhibitory pools. For LDDM (**Figs. 10A-D**), we assumed *ω*_*ij*_ = 1, *τ* _*ij*_ = *τ* _*G*_ = *τ* _*D*_ = 100 m*s*, input scale *S* = 256, decision threshold = 70Hz, and dt = 1 ms. Panel **A** illustrated the temporal dynamic of excitatory pools (R1 and R2) under input coherence of 25% between control (GABAergic weight = 1.0) and agonist (GABAergic weight = 3.8) conditions (other parameters used were *α* = 5, *β* = 1.4, *σ* = 0). Panel **B** examined the predicted RT and choice accuracy over different input coherences (c’ = [0, 3.2, 6.4, 12.8, 25.6, 51.2] %) and levels of GABAergic activities (from 1 (control) to 4 (enhanced)) (*α* = 10, *β* = 1.1, *σ* = 2, and 10000 repetitions). Panel **C** showed the chromomeric and psychometric curves continuous input coherences (1% – 100%) under the section between control (GABAergic weight = 1.0) and GABAergic agonist (GABAergic weight = 1.8). Panel **D** scanned the full parameter space of *α* and *β* between the contrast of control (GABAergic weight = 1.0) and GABAergic agonist (GABAergic weight = 1.8) (c’ = 3.2%, *σ* = 2.0 and 10000 repetitions). For RNM (**Figs. 10E-G**), we used the parameters specified in Wong and Wang, 2006 for the mean-field rate model. GABAergic weight was manipulated by weighting the inhibitory connection in the model. Panel **E** illustrated the noiseless neural dynamics of RNM using the same input coherences and GABAergic enhancement levels as in panel **A**. Panel F was set to compare with panel **B**, thus the input coherences and GABAergic activation kept the same as in panel **B**, with noise amplitude set as *σ* = .02 under the framework of Wong and Wang, 2006. Panel **G** showed the chromomeric and psychometric function predicted by RNM under the same input and GABAergic assumptions as in panel **C**.

## Data availability

Simulation data and empirical data presented in this paper will be available upon publication at https://osf.io/ygr57/?view_only=ea785b6cba18430cbb3d68325bd803e3.

## Code availability

MATLAB code used for simulation and fitting the empirical data will be available upon publication at https://osf.io/ygr57/?view_only=ea785b6cba18430cbb3d68325bd803e3.

