## Supplementary material for "Flexible control of representational dynamics in a disinhibition-based model of decision making"

List of items included in the Supplemental Results

**Motifs tested and compared for normalized coding and winner-take-all choice**

Is the LDDM architecture the only circuit that satisfies the requirements of normalized coding and WTA choice, or can these functions be integrated by an alternative circuit motif? We tested a series of motifs and found local disinhibition is critical for the integration of normalized valuation and choice functions. To do this, we tested four types of modifications that might enhance mutual competition between the option-specific local sub-circuits (**Fig. S1A**): a) *Recurrent self-excitation* (loops weighted by $\alpha$), with self-amplification of each *R* unit, a property shown to be important for mutual competition in the RNM. b) *Local disinhibition* (loops weighted by $\beta$), which is the focus of the main text, mediated through a third type of disinhibitory unit (*I*) to inhibit the gain control *G* unit in the local sub-circuit; disinhibition releases inhibition of the local *R* but maintains inhibition of the lateral *R*. c) *Cross inhibition* (loops weighted by $\eta$), which directly inhibits the lateral *R* units through *I* units to implement mutual inhibition. d) *Lateral gain control boost* (loops weighted by $\gamma$), which increases the lateral *G* through an excitatory unit (*E*) to realize mutual inhibition.

To see which type of modification(s) is/are critical for integrated value normalization and choice, we tested different combinations of these modifications on the original DNM circuit. The full model with all modifications can be described by a set of differential equations:

$$\tau_R \frac{dR_i}{dt} = -R_i + \frac{V_i + \alpha R_i - \eta I_j}{1 + G_i} \tag{S1}$$

$$\tau_G \frac{dG_i}{dt} = -G_i + \omega \sum_{j=1}^{N} R_j + \gamma E_j - \beta I_i \tag{S2}$$

$$\tau_I \frac{dI_i}{dt} = -I_i + R_i \quad \text{(S3)}$$

$$\tau_E \frac{dE_i}{dt} = -E_i + R_i \quad \text{(S4)}$$

where $i = 1, \ldots, N$ designates choice alternatives, each of which receive input $V_i$, and $\tau_R$, $\tau_G$, $\tau_I$, and $\tau_E$ are the time constants for the $R$, $G$, $I$, and $E$ units. The weights $\omega$ represent the coupling strength between excitatory units $R$ and gain control units $G$, the parameters $\alpha$, $\beta$, $\eta$, and $\gamma$ control the active state of recurrent excitation, local disinhibition, cross inhibition, and lateral gain control boost loops, respectively.

The active and inactive states of the four types of loops can be combined into $2^4 = 16$ possible models. Example dynamics are shown in **Fig. S1B** for each type of model. When local disinhibition ($\beta$) is off (left two columns), the model generates WTA dynamics only when cross inhibition ($\eta$) is on. But the maximum activity in the late stage is restricted to a value lower than the phasic peak during the early stage, contradicting empirical findings that the late stage decision threshold is usually higher than activity in the early phasic peak (Churchland et al., 2008; Kiani et al., 2008; Kiani & Shadlen, 2009; Louie et al., 2011; Roitman & Shadlen, 2002; Rorie et al., 2010; Shadlen & Newsome, 2001; Sugrue et al., 2004). This restriction arises because, with only cross inhibition, local option gain control is not released; this release requires local disinhibition. With local disinhibition ($\beta$) on (right two columns), the models generate WTA dynamics with high activity in the late stage to reach the decision threshold. This is robust even without any other modifications (see the panel with $\eta$ and $\gamma$ off), highlighting the role of local disinhibition in generating WTA competition. For the sake of simplicity, we omitted other non-essential modifications and kept only the loop of local disinhibition. Because recurrent excitation is important for persistent activity and exists widely in cortical circuits, we retained it as well. The modified DNM model with local disinhibition and recurrent self-excitation is the primary model (LDDM) characterized in the main text.

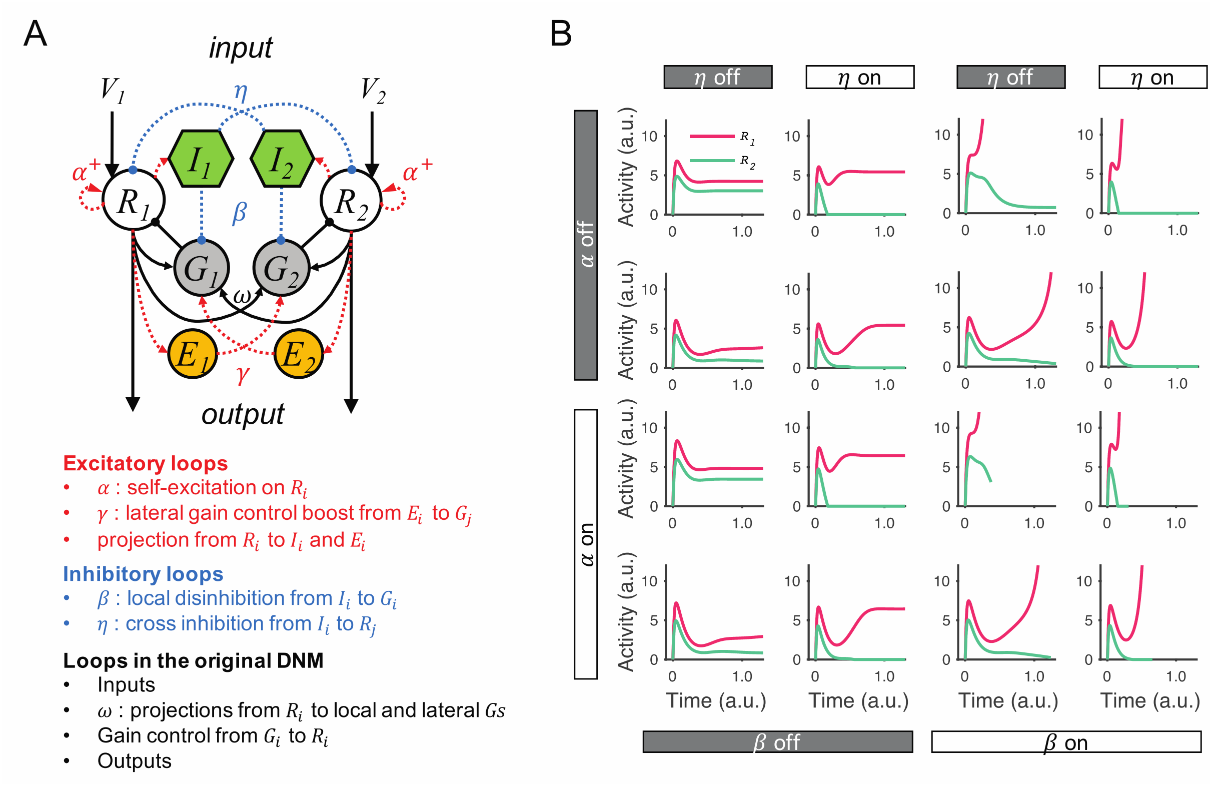

**Figure S1** Testing and comparing different DNM modifications for integrating normalized value coding and WTA competition. **A**. The full model contains all possible modifications that allow the original DNM to generate WTA competition. Modifications: recurrent excitation on $R$ units (controlled by $\alpha$), local disinhibition mediated through $I$ units (controlled by $\beta$), cross inhibition mediated through $I$ units to inhibit the lateral $R$ (controlled by $\eta$), and lateral gain control boost loops mediated through $E$ units (controlled by $\gamma$). **B**. Example $R_1$ and $R_2$ dynamics predicted by the model variants with different combinations of modifications. Four types of modifications result in 16 candidate models. Comparison of the left two columns and right two columns shows that local disinhibition ($\beta$) is required for generating WTA competition and increasing neural activity to a decision threshold.

**Equilibria and stability analysis of the LDDM**

In the main text (**Figs. 3 – 5**), we showed that the LDDM exhibits different pattern of equilibria and stabilities under normalized value coding and WTA competition, mediated through disinhibition. Here we provide detailed mathematical analysis about the equilibria and stability of this dynamic system under different states of disinhibition. We restate here the differential equations of the system (Eq. S5 – S7; the same as Eq. 1 – 3 in the main text):

$$\tau_R \frac{dR_i}{dt} = -R_i + \frac{V_i + \alpha R_i}{1 + G_i} \tag{S5}$$

$$\tau_G \frac{dG_i}{dt} = -G_i + \sum_{j=1}^{N} \omega_{ij} R_j - D_i \tag{S6}$$

$$\tau_D \frac{dD_i}{dt} = -D_i + \beta R_i \tag{S7}$$

where i = 1, …, *N* indicates every input source $V_i$, $\tau_R$, $\tau_G$, and $\tau_D$ are the time constant for the *R*, *G*, and *D* units, $\alpha$ weights the recurrent strength of self-excitation, $\beta$ weights the coupling strength between $R_i$ and disinhibition $D_i$ under external control, $\omega_{ij}$ weights the coupling strength for each gain control unit $G_i$ summing over all of excitatory unit $R_j$. As we assumed the local-option $\omega_{ii}$ and the lateral-option weight $\omega_{ij(i \neq j)}$ are equal, we annotate with the same parameter $\omega$ thereafter.

Equilibria of the system were solved by taking the intersection of the nullclines of all units, i.e., the steady states of each unit. This is obtained by setting $dR_i/dt$, $dG_i/dt$, and $dD_i/dt$ all equal to 0. The solution of the equilibrium state of *R* units ($R_i^*$) can be written as:

$$R_i^* = \frac{V_i}{\omega(\frac{1-\alpha}{\omega} + (1-\frac{\beta}{\omega})R_i^* + \sum_{j\neq i} R_j^*)} \quad \text{(S8)}$$

For a binary input system ($N$ = 2), the six differential equations can be simplified to two equations with only the $R$ units explicitly in the expression (Eq. S9 and S10). Each equation describes the nullcline of a single $R$ unit.

$$\begin{cases} \frac{V_1}{R_1^*} - (\omega-\beta)R_1^* - (1-\alpha) = \omega R_2^* & \text{(S9)} \\ \frac{V_2}{R_2^*} - (\omega-\beta)R_2^* - (1-\alpha) = \omega R_1^* & \text{(S10)} \end{cases}$$

Given that the equilibrium states of the system can be reduced with only R units explicitly in the expression, these equilibrium points can be visualized in the $\mathbb{R}_+^2$ space of $R_1$ and $R_2$ activities as the intersection of the nullclines of the two $R$ units (as shown in **Fig. 3** and **Fig. S2**). The stability of each equilibrium point was then examined by checking the eigenvalues of the Jacobian matrix around it. The equilibrium point is attractive and stable when all of the eigenvalues have negative real parts; the equilibrium point is divergent and unstable when there exist any positive real parts of eigenvalues. By denoting $\boldsymbol{F} = (\boldsymbol{F_{R_1}}, \boldsymbol{F_{G_1}}, \boldsymbol{F_{D_1}}, \boldsymbol{F_{R_2}}, \boldsymbol{F_{G_2}}, \boldsymbol{F_{D_2}})$ as the differential equations for all units in their steady states, the Jacobian matrix around the point can be written as S11:

$$
\boldsymbol{J} = \begin{bmatrix}
\frac{\partial F_{R_1}}{\partial R_1} & \frac{\partial F_{R_1}}{\partial G_1} & \frac{\partial F_{R_1}}{\partial D_1} & \frac{\partial F_{R_1}}{\partial R_2} & \frac{\partial F_{R_1}}{\partial G_2} & \frac{\partial F_{R_1}}{\partial D_2} \\
\frac{\partial F_{G_1}}{\partial R_1} & \frac{\partial F_{G_1}}{\partial G_1} & \frac{\partial F_{G_1}}{\partial D_1} & \frac{\partial F_{G_1}}{\partial R_2} & \frac{\partial F_{G_1}}{\partial G_2} & \frac{\partial F_{G_1}}{\partial D_2} \\
\frac{\partial F_{D_1}}{\partial R_1} & \frac{\partial F_{D_1}}{\partial G_1} & \frac{\partial F_{D_1}}{\partial D_1} & \frac{\partial F_{D_1}}{\partial R_2} & \frac{\partial F_{D_1}}{\partial G_2} & \frac{\partial F_{D_1}}{\partial D_2} \\
\frac{\partial F_{R_2}}{\partial R_1} & \frac{\partial F_{R_2}}{\partial G_1} & \frac{\partial F_{R_2}}{\partial D_1} & \frac{\partial F_{R_2}}{\partial R_2} & \frac{\partial F_{R_2}}{\partial G_2} & \frac{\partial F_{R_2}}{\partial D_2} \\
\frac{\partial F_{G_2}}{\partial R_1} & \frac{\partial F_{G_2}}{\partial G_1} & \frac{\partial F_{G_2}}{\partial D_1} & \frac{\partial F_{G_2}}{\partial R_2} & \frac{\partial F_{G_2}}{\partial G_2} & \frac{\partial F_{G_2}}{\partial D_2} \\
\frac{\partial F_{D_2}}{\partial R_1} & \frac{\partial F_{D_2}}{\partial G_1} & \frac{\partial F_{D_2}}{\partial D_1} & \frac{\partial F_{D_2}}{\partial R_2} & \frac{\partial F_{D_2}}{\partial G_2} & \frac{\partial F_{D_2}}{\partial D_2}
\end{bmatrix}
= \begin{bmatrix}
\begin{matrix} -1 + \frac{\alpha}{1 + G_1^*} & -\frac{V_1 + \alpha R_1^*}{(1 + G_1^*)^2} & 0 \\ \omega & -1 & -1 \\ \beta & 0 & -1 \end{matrix} & \begin{matrix} 0 & 0 & 0 \\ \omega & 0 & 0 \\ 0 & 0 & 0 \end{matrix} \\
\begin{matrix} 0 & 0 & 0 \\ \omega & 0 & 0 \\ 0 & 0 & 0 \end{matrix} & \begin{matrix} -1 + \frac{\alpha}{1 + G_2^*} & -\frac{V_2 + \alpha R_2^*}{(1 + G_2^*)^2} & 0 \\ \omega & -1 & -1 \\ \beta & 0 & -1 \end{matrix}
\end{bmatrix} \quad \text{(S11)}
$$

We examined the configuration of nullclines and checked the eigenvalues of the Jacobian matrix across a wide range of parameter values $\alpha$ and $\beta$, and set $\omega$ as a unit value of 1 for the sake of simplicity. The property of the system under equivalent inputs is a critical test since it determines whether the system is able to implement a WTA choice and select an option. Thus, we examined the property of the system for WTA under equal inputs. Examining the full space of $\alpha$ and $\beta$ revealed five territories distinguished by the number of equilibrium points and their stabilities (**Fig. S2A**). For each territory, the configuration of nullclines are illustrated in **Fig. S2** labeled by color. **Dark green region**: When disinhibition is smaller ($\beta < 1$), $\alpha$ and $\beta$ show a trade-off in generating WTA competition. When both $\alpha$ and $\beta$ are small, the system generates a unique equilibrium point of normalized coding (dark green region in **Fig. S2A**, nullclines shown in **Fig.**

**S2B**). Eigenvalues in this regime show all negative real parts on this equilibrium point, indicating it is a stable equilibrium. **Blue region**: As $\alpha$ values increase (at smaller $\beta$ values), the system generates three equilibrium points (**Fig. S2D**), with two high-contrast (stable) attractors at the peripheral and one (unstable) repellor in the center of space $R_1$-$R_2$. Neural activities of $R_1$ and $R_2$ with equal initial values bifurcate into the high-contrast attractors to realize WTA competition (example traces shown in red and blue lines). **Green region**: When the strength of disinhibition increases ($\beta > 1$), most of the regimes (yellow and red regions) show the properties of WTA competition except for a small regime when $\alpha < 1$ (green region). In the green region, the nullclines of $R_1$ and $R_2$ still intersect on three equilibrium points but, in contrast to the blue region, the two points with high contrast of $R_1$-$R_2$ activities are unstable and the equilibrium point in the center is stable (**Fig. S2C**). The neural activity of $R_1$ and $R_2$ is restricted under a value of $\frac{V_i}{1-\alpha}$, which is lower than the high-contrast equilibria, therefore, the system maintains normalized coding. **Yellow region**: When disinhibition is large ($\beta > 1$), most of the parameter regime in the yellow region shows only one repellor at the center (**Fig. S2E**). The activities of $R_1$ and $R_2$ bifurcate from the center repellor to the high-contrast corners. The restriction of maximum activity depends on the value of $\alpha$. When $\alpha < 1$ , the model predicts limited value of activity on each $R$ unit as ($\frac{V_i}{1-\alpha}$) (vertical and horizontal dashed lines in **Fig. S2E**). When $\alpha \geq 1$, the model predicts no boundary on the maximum activities (though a boundary may still need to be considered because of biological constraints). **Red region**: When disinhibition is extremely large ($\beta > 2$), the two nullclines show no intersections (**Fig. S2F**). Most of the other features in this region are similar to the yellow region. The neural activities of $R_1$ and $R_2$ bifurcate from initial values from the center to the corners of high contrast (example traces shown in red and green thin lines). The boundary of neural activity is predicted when $\alpha < 1$ and not accounted when $\alpha \geq 1$.

Taken together, the five territories can be simplified to two regions based on the properties of the system in implementing either normalized coding or WTA competition as discussed in the main text (**Fig. 5E**). These two regions show clear-cut dichotomous separation in the two-dimensional space of recurrent excitation weight ($\alpha$) and local disinhibition weight ($\beta$).

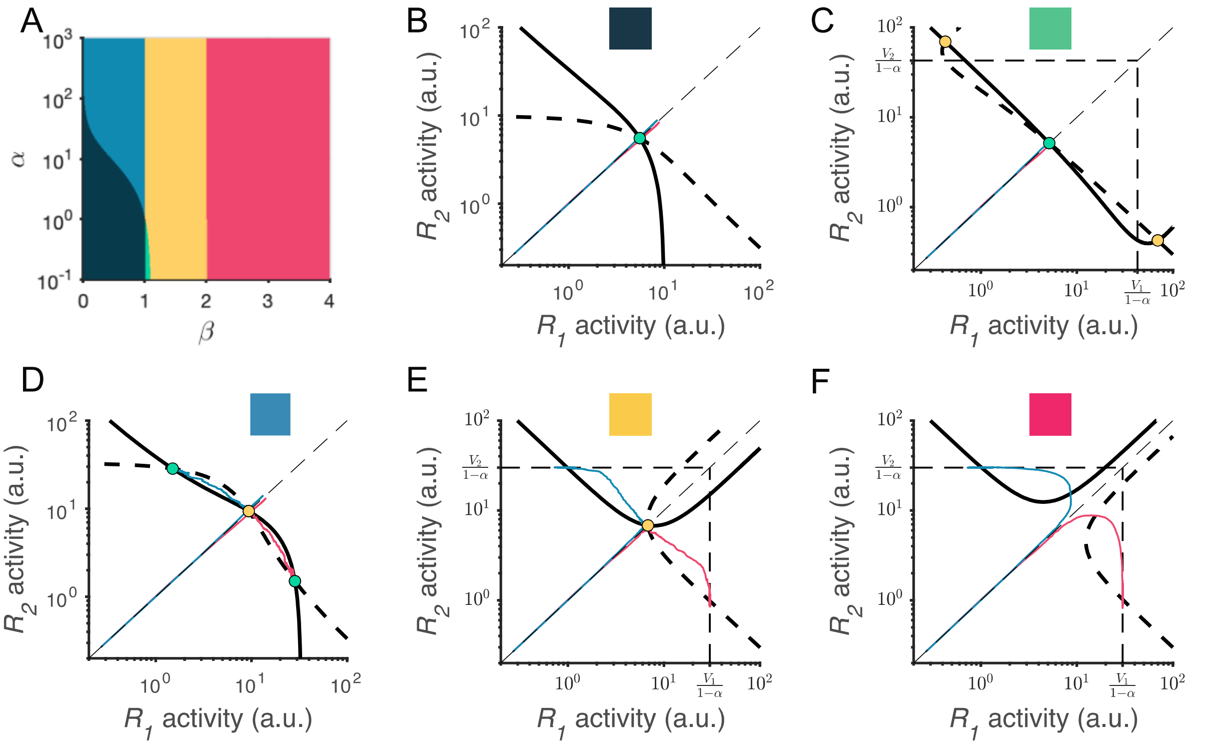

**Figure S2** Phase-plane analyses of the LDDM across a wide range of recurrent excitation strengths ($\alpha$) and local disinhibition strengths ($\beta$). **A**. The five different territories in the space of $\alpha$ and $\beta$ distinguished by the patterns of equilibria and stabilities of the system. **B-F**. Example nullclines of $R_1$ (solid bold line) and $R_2$ (dashed bold line) under each territory of parameter regime indicated by color. Nullclines intersect on equilibrium points, denoted as stable (green dots) or unstable (yellow dots). Example traces of $R_1$ and $R_2$ activities from equal initial values (red and green thin lines) are shown in each panel. The dark green and green regions predict normalized coding attractors but not WTA choice. The blue, yellow, and red regions predict WTA choice but having no normalized coding attractors. The vertical and horizontal dash lines indicate the predicted maximum activities when $\alpha < 1$. Parameters used in these panels are, B: $\alpha = 5, \beta = .3$; C: $\alpha = .3, \beta = 1.01$; D: $\alpha = 10, \beta = .7$; E: $\alpha = 0, \beta = 1.5$; F: $\alpha = 0, \beta = 2.5$. Other parameters apply for all panels: $V_1 = V_2 = 30$, $\omega_{ij} = 1$.

**Fit the RNM to perceptual choice dataset**

In order to compare the model performance in predicting choice behaviors, we fit the original RNM to the classical perceptual decision dataset (Roitman and Shadlen, 2002). We used the reduced form of the RNM (Wong & Wang, 2006). We set eight parameters in the reduced model (see Appendix, Wong & Wang, 2006) as free parameters to fit: self-excitatory coupling weights $JN_{1,1} = JN_{2,2}$, mutual inhibitory coupling weights $JN_{1,2} = JN_{2,1}$, non-selective input $I_0$, noise amplitude of OU process $\sigma_{noise}$, input scale $\mu_0$, synaptic kinetic parameter $\gamma$, initial value $H_0$, and time constant $\tau_S$. The other parameters that describing the input-output relationship of a single cell were set as the same in the paper: *a* = 270 $(\text{VnC})^{-1}$, *b* = 108 Hz, *d* = 0.154 s. The time constant for the AMPA receptor $\tau_{AMPA}$ is fixed as 2 ms. The optimization used is the same as fitting the LDDM (see description of QMLE in **Methods**). Time step dt was set as .001 s.

In the best fitting results, most of the best-fitting parameters are close to the values selected in the original paper (Wong & Wang, 2006) ($JN_{1,1} = JN_{2,2} = .2632, JN_{1,2} = JN_{2,1} = .0224, I_0 = .2647$, $\sigma_{noise} = .0709$, $\mu_0 = 55.63$, $\gamma = .5887$, $H_0 = 2.622$, and $\tau_S = .1672$). The RNM captures well the shape of RT distribution. The fitting performance was shown as histogram of RT distributions (**Fig. S3A**) and a Q-Q plot of RT quantiles as functions of chosen proportion (**Fig. S3B**), as well as aggregated values based on input coherences (**Fig. S3C**) (negative log-likelihood = 16587). The predicted dynamics generated using the best-fit parameters to behavior show a slightly different pattern than the empirical data (**Fig. S3D**). Sorting to the onset of stimuli, the model predicted ramping-up speeds increase with input values. Chosen activity shows an increase with input coherence at a rate of 4.97 spikes/second/100% coherence (**Fig. S3E line a**); unchosen activity shows a decrease of 4.58 spikes/second/100% coherence (**Fig. S3E line b**). Whereas it didn't show as a pattern of competition since the best-fit mutual inhibition parameter equals to zero. Sorting to the time of action execution, the chosen signals hit the same decision threshold regardless of input strength (**Fig. S3F line c,** difference across coherence conditions: .0356 spikes per second per 100% coherence) and overshot because of the attraction property of the network under self-excitation. The unchosen side remains a graded coding of input strength (**Fig. S3F line d,** -7.98 spikes/second/100% coherence).

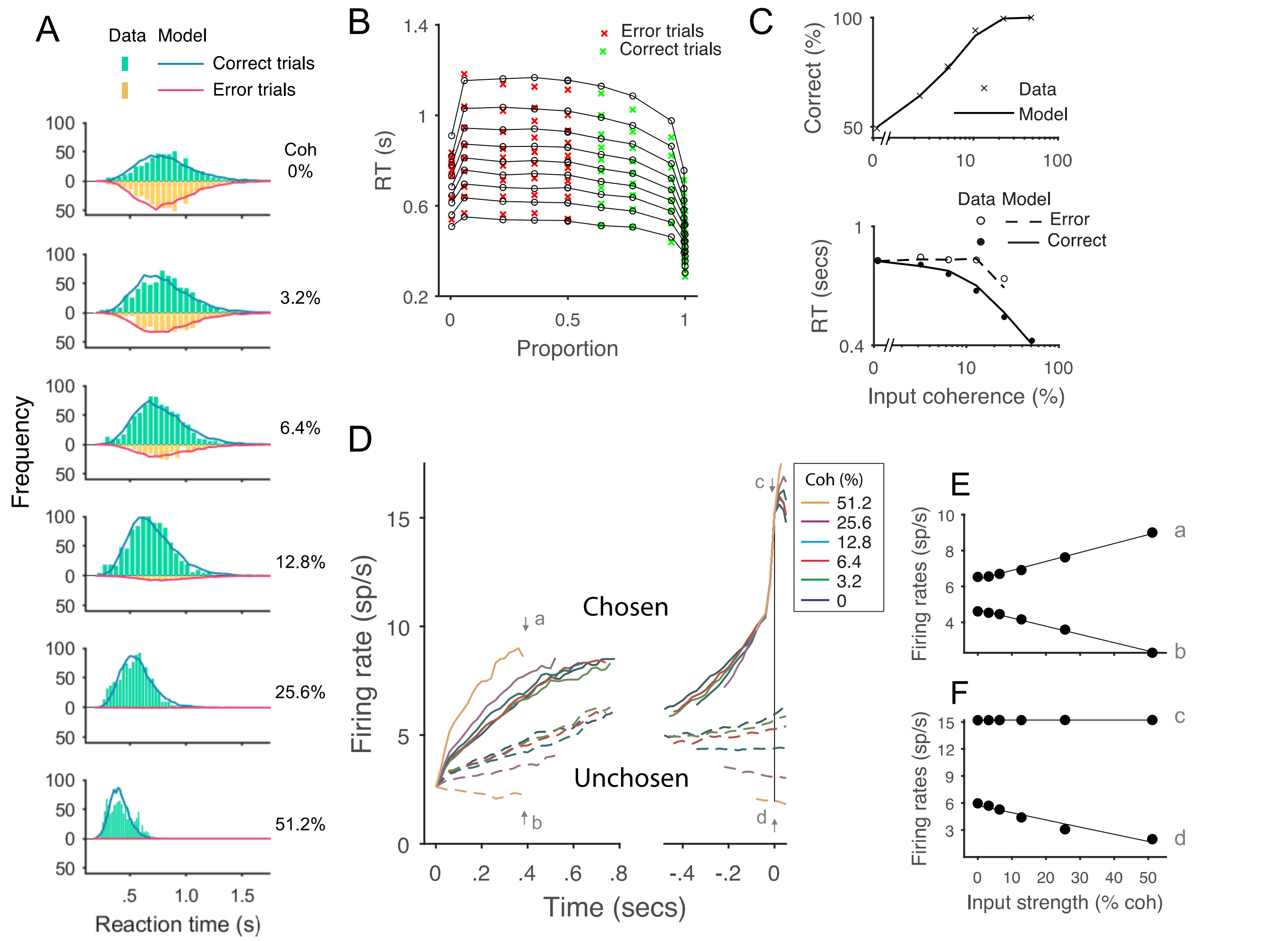

**Figure S3.** Fit the original RNM to the classical dataset (Roitman and Shadlen, 2002). **A.** Model predicts RT distributions (lines) match the histogram of empirical RT distribution (bars), with correct and error trials separated (indicated by color) across levels of input strength (% coherence). **B.** Re-plot the fitting results in a quantile-quantile (Q-Q) plot, with nine quantiles of RT from each condition stacked on the choice proportion of the condition (0 - .5 are error trials, shown in red cross; .5 - 1 are correct trials, shown in green cross). Model predicts well the choice proportion and the shape of RT distribution. **C.** Model predicted psychometric function and chronometric function aggregated input strength. Upper panel: Choice accuracy (lines) fit well to the empirical data (cross). Lower panel: The predicted RT for correct (solid line) and error (dashed line) trials captures well the RT for correct (filled dots) and error (empty dots) trials in empirical data. **D.** The aggregated neural dynamics from the best-fit model of RNM. Left, mean-field activities on the excitatory pools aligned to the onset of stimulus inputs. The best fitted mutual inhibition parameter is 0. The ramping-up speeds differ over input strengths (see detailed pattern in **E**). Whereas the dynamics didn't show competitions between the chosen (solid lines) and unchosen (dashed lines) signals. Right, mean-field activities on the excitatory pools aligned to the time of choice execution. The chosen signals overshot the threshold 15Hz because of the self-excitation in the circuits. The unchosen signals show graded coding of the input strengths (see detailed pattern in **F**). **E.** Effect of input strength on early stage model activity shortly after stimulus onset, at times indicated by arrows a (chosen side) and b (unchosen side). The activities linearly increase and decrease with input strength but exhibit very subtle competition. **F.** Effect of input strength on late stage model activity on the time point of choice, indicated by arrows c and d. Chosen unit activity reaches a common decision bound and show little input dependence, while unchosen activity remains at lower levels, graded coding of input strengths.

**Fit the RNM to normalized value coding dataset**

In order to quantify the performance of the RNM in predicting normalized value coding, we fit the reduced form of RNM (Wong and Wang, 2002) with 4 free parameters ($JN_{i,i,i}$, $JN_{i,j,k(i \neq j \neq k)}$, $I_0$, and a scaling parameter $S$ applied to the predicted neural firing rates) to a normalized value coding dataset (**Fig. 4** in Louie et al., 2011). Other parameters are set the same as reported in the original paper (Wong and Wang, 2002), except that the noise term $\sigma$ is set as zero. The RNM is expanded to a trinary choice circuit, with three selective populations wired together based on the same rules specified in the original paper (Wong and Wang, 2002). We study the predicted neural activity on the pool 1 that receiving direct input from $V_1$ and too investigate how the activity of pool 1 changes with the values of contextual inputs $V_2$ and $V_3$. Bayesian adaptive direct search algorithm (BADS, Acerbi & Ma, 2017) is used to minimize the ordinary squared error between the predicted neural firing rates of pool 1 and the empirical neural firing rates data reported in the Fig. 4 of Louie et al., 2011. The best fitting result show that the RNM explains 89.2% of the variance, much worse than the DNM and LDDM we reported in the main text (Best fitting parameters: $JN_{i,i,i}$ = .0055, $JN_{i,j,k(i \neq j \neq k)}$ = .0861, $I_0$ = .3511, $S$ = 1.074). The fitted curve of pool 1 activities illustrated in **Fig. S4**. The RNM fails to capture the intercept of each curve given different $V_{in}$ under the lateral inhibition of $V_{out}$. In addition, the fitted curve across the contextual inputs ($V_{out}$) show linear type of suppression instead of a curvature shows in the LDDM and DNM, which matches the principle of divisive normalization (**Fig. 4B**). Moreover, after check the non-scaled neural firing rates from the best fitting, we realize that the RNM predict unrealistically low level of activities, with a maximum value ~ 3.5 Hz (**Fig. S4B**), inconsistent with empirical findings.

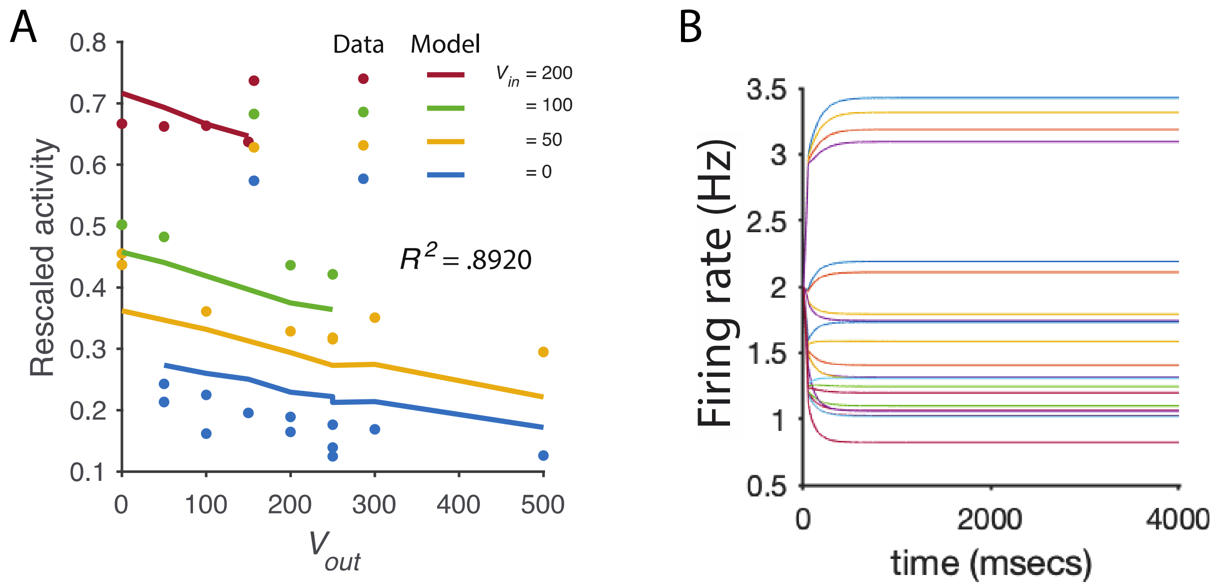

**Figure S4. Fit the RNM to the dynamic of normalized value coding. A**. The best fitted neural activity on pool 1 (lines) to the empirical data of the neural activity in the response field getting direct input from $V_{in}$ (points). $V_{in}$ and $V_{out}$ vary across conditions, with each dot indicating a input condition. The RNM does worse than the LDDM and DNM in capturing the intercept on the neural activity across $V_{in}$ values under the modulation of contextual input $V_{out}$, and predicted linear type of lateral inhibition instead of divisive normalization. **B**. The predicted dynamic of neural firing rates without scaling, including the activities of all three pools across different input conditions. The predicted dynamic shows an unrealistic low activity level, inconsistent with empirical observations.

**Analysis for persistent activity**

In the main text, we showed that the LDDM with recurrent excitation predicts persistent activity that maintains input information during delay intervals. Here we provide mathematical analysis of the LDDM differential equations to examine the properties and genesis of this persistent activity. In addition to examining the property of the system with symmetric gain control weights ($\omega_{ii} = \omega_{ij(i\neq j)}$), we expanded our analysis to allow the gain-control weights to be asymmetric; this allows us to examine the robustness of LDDM properties to asymmetric weights.

Equilibrium states of the differential equations (Eq. S5 – S7) after withdrawal of inputs were considered. The gain control weighs $\omega_{ij}$ were split into two parts, with the local-option weight denoted as $w$ ($\omega_{ii} = w$) and the cross-option weight denoted as $v$ ($\omega_{ij(i\neq j)} = v$). The input values were set to zero and local disinhibition was assumed inactive ($\beta = 0$). Equilibria of the system were solved by taking the intersection of the steady states of all units, i.e., when $dR_i/dt$, $dG_i/dt$, and $dI_i/dt$ all equal to 0. When the input terms are set to zero, the solution degrades from Eq. S8 to Eq. S12 as a linear form,

$$wR_i^* + v\sum_{i\neq j} R_j^* = \alpha - 1 \tag{S12}$$

For a binary choice system, the solution of Eq. S12 is denoted in linear algebra as:

$$\begin{bmatrix} w & v \\ v & w \end{bmatrix} \begin{bmatrix} R_1^* \\ R_2^* \end{bmatrix} = \begin{bmatrix} \alpha - 1 \\ \alpha - 1 \end{bmatrix} \tag{S13}$$

The solutions of the equations depend on the value of $\alpha$. When $\alpha \leq 1$, the equations do not provide a positive solution. This explains why the system without recurrent excitation ($\alpha = 0$) cannot generate persistent activity. When $\alpha > 1$, the equations provide positive solutions. The model generates persistent activities in three different patterns depending on the symmetry of gain control weights, i.e., $v < w$, $v = w$, and $v > w$.

First, by assuming $v = w$ and $\alpha > 1$, the nullclines of $R_1$ and $R_2$ overlap on a line of attraction, as shown in the main text (**Fig. S5B**, the same as **Fig. 8B**). Any position on this line is an equilibrium point. This is a special case where the eigenvalues on each point have a real part of zero, therefore, linearization around the equilibrium points cannot tell us their stability. Thus, we checked instead the instantaneous change direction of neural activities across a wide range of initial values to see whether the system converges to the line of attraction. From the differential equations (Eq. S5 – S7), the ratio $dR_1/dR_2$ of the instantaneous change rates of $R_1$ ($dR_1/dt = R_1(1 - \frac{\alpha}{1+G_1})$) and $R_2$ ($dR_2/dt = R_2(1 - \frac{\alpha}{1+G_2})$) keeps the same ratio as the ratio of original activities ($R_1/R_2$), given $G_1 = G_2$ under the assumption of symmetric gain control weights. As a result, for any given initial values, $R_1$ and $R_2$ activities change in the direction that preserves the original ratio until reaching equilibrium on the line of attraction. The instantaneous changes of $R_1$ and $R_2$ are shown as a vector field (red arrows) in **Fig. S5B**. Thus, any positive initial values will drop into an equilibrium state with the ratio of $R_1^*/R_2^*$ maintaining the ratio of initial values, which preserves the ratio of inputs when the activities are inherited from the stage of value representation. **Fig. S5E** shows example dynamics of $R_1$ and $R_2$ under different ratios of input values (**Fig. S5G**). The activities show the characteristic dynamic of divisive normalization during the inputs and preserve this input information after withdrawal of inputs.

However, since the values of $R_1^*$ and $R_2^*$ are complementary on the line of attraction, any combinations of values with a constant sum satisfies the equilibrium. Thus, any disturbance to the system (e.g. random noise) will drive $R_1^*$ and $R_2^*$ to deviate from their original ratio resulting in a loss of the coded information about the inputs. Noise-driven drift on the line of attraction will cause decaying of the coded value information over time, consistent with the degradation attribute of working memory (Barrouillet et al., 2011; Barrouillet & Camos, 2012; Lee & Harris, 1996; Paivio & Bleasdale, 1974; Portrat et al., 2008).

In addition, under the special condition of symmetric gain control weights ($v = w$), the formula in Eq. S13 can be easily expanded to multiple inputs with the equilibrium delay interval activities defined by:

$$\sum_{i}^{N} R_i^* = \frac{\alpha - 1}{w} \tag{S14}$$

The summed value of all $R$ units equals to a constant $\frac{\alpha-1}{w}$. When the number of inputs ($N$) increases, the activity shared by each $R$ unit decreases and leads to a lower signal relative to noise scale. Thus, as the number of coded items increasing, the information kept during persistent activity may become less accurate considering lower signal-to-noise ratio. This may explain another important attribute of working memory – the constraint of working memory span (Cowan, 2010, 2016; Engle, 2001, 2002; Oberauer et al., 2016).

Second, by assuming $v < w$ and $\alpha > 1$, the nullclines of $R_1$ and $R_2$ intersect on a unique equilibrium point, where $R_1$ and $R_2$ share the same value $\frac{\alpha-1}{w+v}$ (**Fig. S5A**). The point is confirmed as attractive by linearization. Any positive initial values on the space of $R_1$ and $R_2$ will converge into this point, which is visualized in the instantaneous change ranges of $R_1$ and $R_2$ (red arrows) for a wide range of given initial values (**Fig. S5A**). Thus, $R_1$ and $R_2$ will gradually converge to be equal and the original information about input values will be lost. Nevertheless, the dynamic of information losing is based on the level of asymmetry of $\omega_{ij}$. For a close-to-symmetric $\omega_{ij}$ matrix, the input information can be still preserved for a considerable amount of time. We showed example dynamics of information loss in **Fig. S5D** (input values shown in **Fig. S5G**). After withdrawal of inputs, the $R$ unit activities collapse into the same level and the coded ratio information gradually diminishes (simulation parameters: $\alpha = 10, w = 1, v = .7, \beta = 0$).

Finally, by assuming $v > w$ and $\alpha > 1$, the nullclines of $R_1$ and $R_2$ intersect on a unique equilibrium point, which is confirmed as unstable by linearization (**Fig. S5C**). Any initial values of activities on the space will diverge into the upper-left or bottom-right corner of the space generating high contrast between $R_1$ and $R_2$, with the higher activity as $\frac{\alpha-1}{w}$ and the lower activity suppressed to zero. The instantaneous change rates of $R_1$ and $R_2$ (red arrows) are visualized in the vector field in **Fig. S5C**. The instantaneous change direction bifurcates at the line of $R_1 = R_2$, biased to the side associated with higher initial activity. As an outcome, the $R$ unit with higher initial values tends to increase while the opponent unit tends to be suppressed to zero, a process that implements WTA competition before the action stage but with constrained higher activity. Example $R_1$ and $R_2$ activity dynamics are shown in **Fig. S5F** (input values shown in **Fig. S5G**). After withdrawal of inputs, $R_1$ activities with different preceding input values collapse onto the same level of high activity, while $R_2$ activities with lower input values are suppressed to zero. Thus, the system gradually switches from the normalized coding of inputs to a categorical coding of choice over the delay interval.

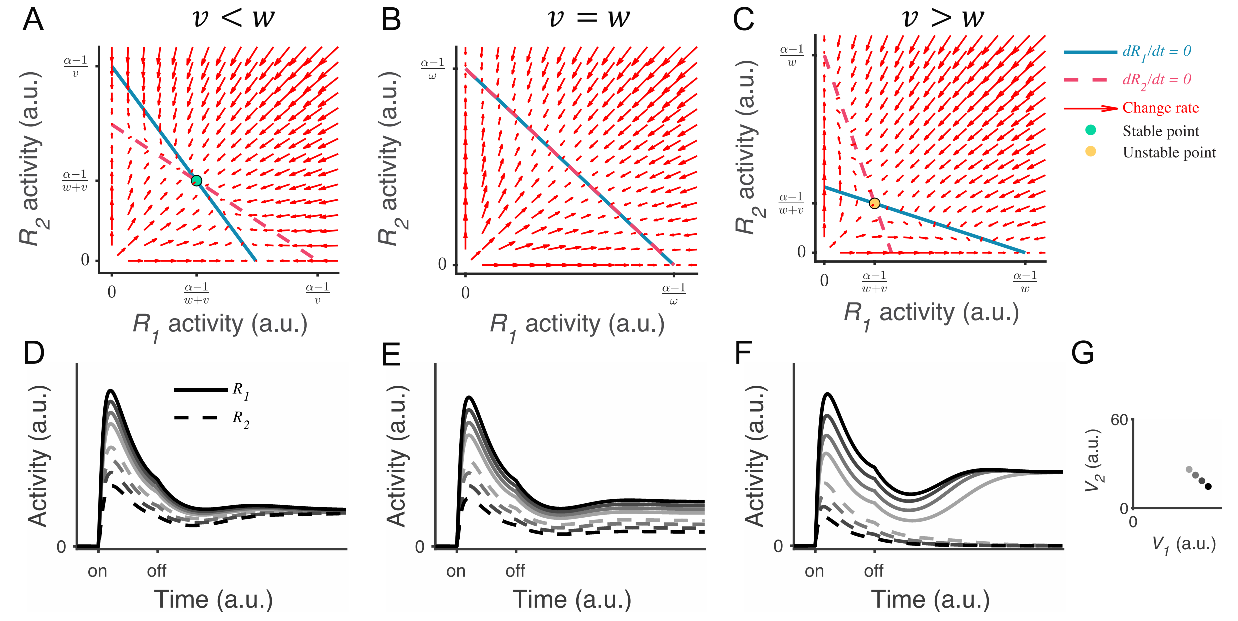

**Figure S5** Analysis of persistent activity under generalized gain control weights. **A**-**C**. Phase-plane analysis shows that systems with different gain control weights have different patterns of equilibria and stabilities. **A**. When the lateral gain control ($v$) is weaker than the local gain control ($w$), the nullclines of $R_1$ (blue solid) and $R_2$ (red dashed) intersect on an attractive unique equilibrium point. Vector filed (red arrows) indicates the instantaneous change rate of $R_1$ and $R_2$ at given initial values. Any initial values converge into the equilibrium point, with $R_1$ and $R_2$ sharing the same value $\frac{\alpha-1}{w+v}$. **B**. When $v = w$, the nullclines of $R_1$ and $R_2$ overlap on the line of attraction. Vector field shows that any initial values converge onto the line of attraction along the direction that preserves the original input ratio. **C**. When $v > w$, the nullclines of $R_1$ and $R_2$ intersect on a unique but unstable point. Any initial values diverge from the point and bias to the side with higher initial value, realizing WTA competition. **D**-**F**. Example neural dynamic on $R_1$ and $R_2$ when under different input ratios (indicated by grayscale and shown in **G**). Corresponding to the phase-plane analysis in **A**-**C**, the activities of $R_1$ and $R_2$ gradually converge onto the same value when $v < w$, keep the input ratio when $v = w$, and diverge based on the input ratio when $v > w$. **G**. Input values used in the [illegible]

**Persistent activity under local disinhibition**

Here, we examine whether persistent activity can also exist with active local disinhibition. We showed in the main text that persistent activity in the working-memory task switches to WTA choice under the dynamic control of disinhibition (**Fig. 8D-F**). How does the transition from persistent activity to WTA choice happen? Furthermore, how might disinhibition change the dynamic pattern of persistent activity during a delay interval?

The analysis was based on the differential equations of the system with symmetric gain control weights and without inputs (Eq. S5 – S7). The equilibrium solution is given by:

$$(\omega - \beta)R_i^* + \omega \sum_{i \neq j} R_j^* = \alpha - 1 \qquad \text{(S15)}$$

With binary inputs, the solution can be thus written as:

$$\begin{bmatrix} \omega - \beta & \omega \\ \omega & \omega - \beta \end{bmatrix} \begin{bmatrix} R_1^* \\ R_2^* \end{bmatrix} = \begin{bmatrix} \alpha - 1 \\ \alpha - 1 \end{bmatrix} \qquad \text{(S16)}$$

Beside the impact of recurrent excitation ($\alpha$) discussed above, equilibrium responses are determined by the relative strength between disinhibition ($\beta$) and the gain control weight ($\omega$). We examined three separate conditions: $\beta = 0$, $0 < \beta < \omega$, and $\beta > \omega$. We have already shown the analysis for the special case when $\beta = 0$ above (phase plane analysis and example dynamic shown in **Fig. S5B**) and replotted in **Fig. S6A** for the sake of comparison with other two conditions.

By assuming $0 < \beta < \omega$, the nullclines of $R_1$ and $R_2$ intersect on a unique equilibrium point, whose stability was confirmed as unstable after checking the eigenvalues of Jacobian matrix around the point (**Fig. S6B**). Any initial values on the space will diverge into the upper-left or bottom-right corner of the space, with the higher activity value as $\frac{\alpha-1}{\omega-\beta}$, and the lower activity value as zero. We show the instantaneous change rates of $R_1$ and $R_2$ at given initial values in the vector field (red arrows) (**Fig. S6B**). In **Fig. S6E**, we show example $R_1$ and $R_2$ activity dynamics (value setting kept the same as in **Fig. S5G**). All of the $R_1$ with larger input values converge into the same level of activity after withdrawal of inputs, while all of the $R_2$ with lower input values are suppressed to zero, implementing a WTA competition. Thus, the system gradually switches from normalized coding of input values to categorical choice from the early to the late stage of persistent activity.

By assuming $\beta > \omega$, most of the features are similar to the previous situation, except that the model now predicts no constraints on the maximum activity (**Fig. S6C**). The system shows nullclines with an intersection at a unique repellor. The activities of $R_1$ and $R_2$ bifurcate at the line of $R_1 = R_2$. The example dynamics show that the activity of $R_1$, which has higher initial value, increases to an unlimited level and thus will reach a decision threshold. The rising speed of $R_1$ depends on the advantage of $R_1$ over $R_2$ as defined by their initial values.

Taken together, these results show that persistent activity is present as normalized coding of input values only with symmetric gain control weights ($w = v$) and inactive disinhibition ($\beta$). When disinhibition has a moderate strength ($0 < \beta < \omega$), the persistent activity gradually transitions from value coding to categorical choice coding but avoids hitting the decision threshold. When disinhibition is strong enough ($\beta > \omega$), the system generates WTA competition and reaches the decision threshold.

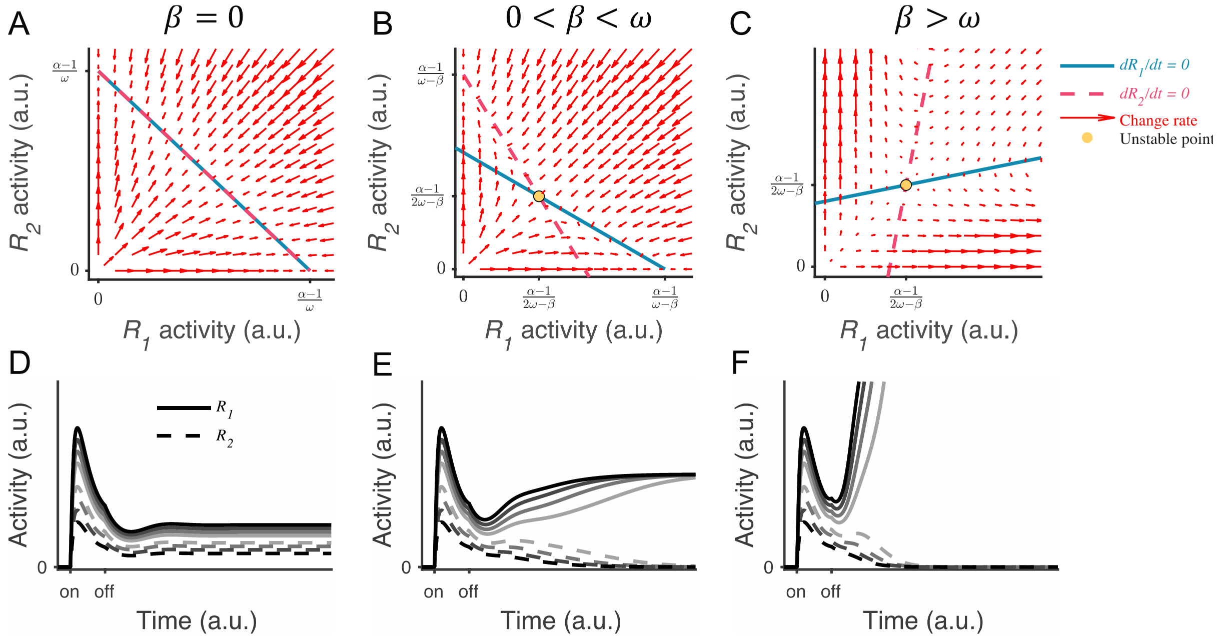

**Figure S6** LDDM persistent activity under different levels of local disinhibition. **A-C**. Phase-plane analysis of persistent activity for the situations of inactive disinhibition ($\beta = 0$, **A**), moderate intensity of disinhibition ($0 < \beta < \omega$, **B**), and strong disinhibition ($\beta > \omega$, **C**). When $\beta = 0$, the nullclines of $R_1$ (blue solid) and $R_2$ (red dashed) intersect on a line of attraction, resulting in normalized value coding. When $0 < \beta < \omega$ and $\beta > \omega$, the $R_1$ and $R_2$ nullclines intersect on an unstable repellor. The vector field (red arrows) shows the instantaneous change rate of $R_1$ and $R_2$ at given initial values. **D-F**. Example $R_1$ and $R_2$ dynamics under different input values (indicated by grayscale in **Figure S5G**). When $\beta = 0$ (**D**), the activities of $R_1$ (solid) and $R_2$ (dashed) maintain the normalized coding of input values during persistent activity. When $0 < \beta < \omega$ (**E**), $R_1$ and $R_2$ gradually transition from coding of the normalized value to coding of categorical choice but the activity is still beneath the decision threshold. When $\beta > \omega$ (**F**), $R_1$ and $R_2$ exhibit WTA dynamics and the winner [illegible]
